# Finite Sample Corrections for Average Equivalence Testing

**DOI:** 10.1101/2023.03.11.532179

**Authors:** Younes Boulaguiem, Julie Quartier, Maria Lapteva, Yogeshvar N Kalia, Maria-Pia Victoria-Feser, Stéphane Guerrier, Dominique-Laurent Couturier

## Abstract

Average (bio)equivalence tests are used to assess if a parameter, like the mean difference in treatment response between two conditions for example, lies within a given equivalence interval, hence allowing to conclude that the conditions have ‘equivalent’ means. The *Two One-Sided Tests* (TOST) procedure, consisting in testing whether the target parameter is respectively significantly greater and lower than some pre-defined lower and upper equivalence limits, is typically used in this context, usually by checking whether the confidence interval for the target parameter lies within these limits. This intuitive and visual procedure is however known to be conservative, especially in the case of highly variable drugs, where it shows a rapid power loss, often reaching zero, hence making it impossible to conclude for equivalence when it is actually true. Here, we propose a finite sample correction of the TOST procedure, the *α*-TOST, which consists in a correction of the significance level of the TOST allowing to guarantee a test size (or type-I error rate) of *α*. This new procedure essentially corresponds to a finite sample and variability correction of the TOST procedure. We show that this procedure is uniformly more powerful than the TOST, easy to compute, and that its operating characteristics outperform the ones of its competitors. A case study about econazole nitrate deposition in porcine skin is used to illustrate the benefits of the proposed method and its advantages compared to other available procedures.

## 1 Introduction

Equivalence tests, also known as similarity or parity tests, have gained significant attention during the past two decades. They originated from the field of pharmacokinetics, ^1,2^where they are called bioequivalence tests and have numerous applications in both research and production. ^3^ They find their most common application in the manufacturing of generic medicinal drugs, where, by proving that the generic version has a similar bioavailability to its well-studied brand-name counterpart, the manufacturer can considerably shorten the approval process for the generic drug. ^4^ Moreover, equivalence tests have attracted growing interest in other domains and for other types of purposes, such as in production when, for example, the mode of administration is altered or when the production site is changed, ^5^ or in the social and behavioral sciences for the evaluation of replication results and corroborating risky predictions. ^6^ Very recent literature reflects the expanding use of equivalence tests across a growing range of domains. Examples include the investigation of the equivalence of virtual reality imaging measurements by feature, ^7^ of cardiovascular responses to stimuli by sex, ^8^ of children neurodevelopment, ^9^ chemotherapy efficacy and safety by treatment, ^10^ of post-stroke functional connectivity patterns by patient group, ^11^ of risk-taking choices by moral type, ^12^ and of 2020 US presidential election turnout by political advertising condition. ^13^ Review articles have also appeared, for example, in food sciences, ^14^ in psychology, ^15^ in sport sciences, ^16^ and in pharmaceutical sciences. ^17^

Equivalence testing implies defining an equivalence region within which the parameter of interest, such as the difference between outcome means measured under two conditions, would lie, for these conditions to be considered equivalent. Indeed, when comparing two treatments, for example, differences in therapeutic effects that belong to the equivalence region would typically be considered as negligible or irrelevant. This is different from standard equality-of-means hypothesis tests in which the null and the alternative hypothesis are interchanged and the null hypothesis states that both means are equal rather than equivalent.

Formally, a canonical form for the average equivalence problem consists of two independent random variables 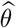 and 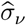 having the distributions

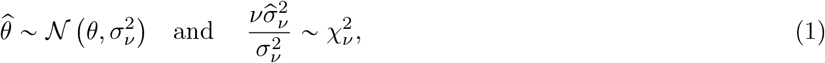

where *θ* and 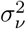 respectively denote the target equivalence parameter and its variance, depending on the number of the degrees of freedom *ν* which is a function of the sample size and total number of parameters. This setting is very general. It covers cases where, for example, the bioequivalence parameter corresponds to the difference in means of the (logarithm of pharmacokinetic) responses between two experimental conditions, or to an element of the parameter vector of a (generalised) linear mixed effect model, like the difference between the slopes of two conditions in a longitudinal study. The hypotheses of interest are given by

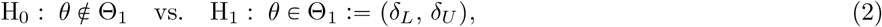

where *δ*_*L*_ and *δ*_*U*_ are known constants. Without loss of generality, it can be assumed that the equivalence limits are symmetrical around zero. In this case, *c* := *δ*_*U*_ = −*δ*_*L*_ so that Θ_1_ =(−*c, c*). Equivalence is typically investigated via the *Two One-Sided Tests* (TOST) procedure, ^18^ consisting in testing whether the target parameter is respectively significantly greater than −*c* and lower than *c*, with the test size, or type-I error rate, controlled at the significance (or nominal) level *α*, usually chosen as 5%. ^19^ More precisely, the TOST is level-*α*, meaning that its size is smaller or equal to *α* (see (4) in Section 2.1, for its formal definition). The most common way of assessing equivalence is to use the *Interval Inclusion Principle* (IIP) and check whether the 100(1 -2*α*)% Confidence Interval (CI) for the target parameter falls within the equivalence margins (−*c, c*). ^3,20^ This strategy has been shown to lead to the same test decision as the TOST procedure if the CI is equi-tailed. ^21,22^

However, it is well known that this procedure can be conservative as the size of the TOST can be considerably lower than the (specified) significance level *α*. This induces a drop in power and therefore to a lower probability of detecting truly equivalent mean effects as such. This problem is particularly noticeable in cases where *σ*_*ν*_ is relatively large. Such situations may occur, for example, when the sample size is determined using an underestimated standard deviation value obtained from a prior experiment, or with studies involving highly variable drugs and in which the sample size that would be needed to achieve reasonable values for *σ*_*ν*_ is unrealistic. For that purpose, Anderson and Hauck ^23^ proposed a test that has greater power than the TOST for situations where *σ*_*ν*_ is relatively large. This test, referred to here as the AH-test, can be liberal and therefore doesn’t control the size. ^22^ In some cases, it can also lead to the equivalence declaration (i.e. acceptance of equivalence through the rejection of the null hypothesis in (2) at the *α* level) when *θ*, the target parameter of interest, falls outside the equivalence interval. ^18^ Brown et al. ^24^ constructed an unbiased test that is uniformly more powerful than the TOST, however, it is computationally intensive and its rejection region may exhibit rather irregular shapes in some cases. ^22^ Berger et al. ^22^ therefore proposed a smoothed version. These tests cannot be assessed using the IIP and the last two are difficult to interpret due to the use of polar coordinates. ^25^

In the specific context of average bioequivalence testing in replicated crossover designs ^26^ for highly variable drugs, i.e., for cases with relatively large *σ*_*ν*_, regulatory authorities have recommended an alternative approach based on the linear scaling of the bioequivalence limits according to the value of the standard deviation within the reference group, called Scaled Average BioEquivalence (SABE), ^27^ also referred to as Average BioEquivalence with Expanding limits (ABEL) in some references, ^26,28,29^ with the constraint that 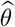 lies within the bioequivalence margins (−*c, c*). These recommendations were issued by the European Medicines Agency (EMA) and the US Food and Drug Administration (FDA). ^20,30^ The amount of expansion is limited by the authorities, and several recent publications have shown that the size of the SABE can be larger than the significance level *α* ^20,26,31,32,33^ and therefore have proposed different ways to correct for it. ^28,34,35,36^ These corrections ensure that the size is smaller than or equal to *α* and lead to acceptance regions that change more smoothly with *σ*_*ν*_ .

In this paper, as an alternative to previous methods, we propose a finite sample correction of the TOST procedure that simply consists in a correction of the TOST’s significance level to guarantee a size-*α* test when *σ*_*ν*_ is known. This correction is design-agnostic and can be used with parallel or (replicated) crossover designs, for example. The corrected significance level *α*^*^ is straightforward to compute and allows to define 100(1 - 2*α*^*^)% CIs used in the classical TOST. Hence, the *α*-TOST essentially corresponds to a finite sample continuous variability correction of the TOST procedure, that leads to an increased probability of declaring equivalence when it is true for large values of *σ*_*ν*_ while maintaining a size of exactly *α* when *σ*_*ν*_ is known. Indeed, the *α*-TOST is shown to be uniformly more powerful than the TOST and, for small to moderate values of *σ*_*ν*_, to be nearly equivalent to the TOST with a comparable power as *α*^*^ ≈ *α* in such cases. Since, in practice, *σ*_*ν*_ needs to be estimated from the data, a straightforward estimator for *α*^*^ is also proposed. It is shown, through an extensive simulation study considering the canonical form defined in (1) and therefore valid in a wide range of settings, that the estimator remains level-*α* and its size stays close to *α*. Our simulation study also considers a version of the TOST that adjusts the equivalence limits *δ*_*L*_ and *δ*_*U*_ instead of the level, to guarantee a size-*α* test and therefore referred to as the *δ* −TOST. Our results show that the *α*−TOST is both more powerful and accurate than the standard TOST and *δ*-TOST, indicating that, when looking for a design-agnostic correction valid in general settings, a correction on the level (*α*-TOST) leads to better operating characteristics. A comparison of the performance of these methods to the corrected SABE, that consists in an adjustment on both the equivalence bounds and the level, is presented in Appendix E in a simple paired setting. More adequate and extensive comparisons, considering the different adjustments proposed by regulatory agencies and other authors, including variants such as the corrected SABE, are needed in the specific case of average equivalence testing with replicated crossover designs and are left for further research.

The paper is organized as follows. The *α*-TOST is presented in Section 2 from a suitable formulation of the TOST. Its statistical properties as well as a simple algorithm to compute *α*^*^ are also provided. In Section 3, an extensive simulation study is used to compare the empirical performances of the *α*-TOST, *δ*-TOST and standard TOST. In Section 4, we consider a case study for which we apply the TOST and the *α*-TOST, as well as other available methods, in order to showcase the advantages of our proposed design-agnostic approach. Finally, Section 5 discusses some potential extensions.

## 2 Equivalence Testing

In this section, we present the methodology for deriving a corrected statistical equivalence test. We first present the TOST and its properties. We then define the *α*-TOST procedure through a natural correction of the TOST, derive its statistical properties, propose an iterative procedure to compute the corrected level *α*^*^ and show that this procedure is exponentially fast. We also show that the *α*-TOST is uniformly more powerful than the TOST.

### 2.1 The TOST Procedure

For testing the hypotheses in (2), the TOST uses the two following test statistics:

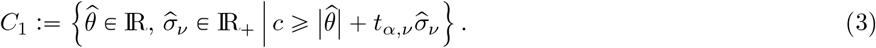

where *Z*_*L*_ tests for H_01_ : *θ* ⩽ −*c* versus H_11_ : *θ* >−*c*, and *Z*_*U*_ tests for H_02_ : *θ* < *c* versus H_12_ : *θ* ⩾ *c*. At a significance level *α*, the TOST therefore rejects H_0_ :=H_01_ ∪ H_02_ (i.e., *θ* ∉ Θ_1_) in favour of H_1_ :” H_11_ ∩ H_12_ (i.e., *θ* ∈ Θ_1_) if both tests simultaneously reject their marginal null hypotheses, that is, if

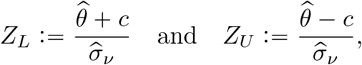

where *t*_*α,ν*_ denotes the upper *α* quantile of a *t*-distribution with *ν* degrees of freedom. The corresponding rejection region of the TOST is given by

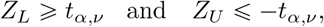

Consequently, equivalence cannot be declared with the TOST for all 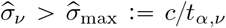_*ν*_, even for 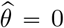 (see also Figure 6 of Section 4).

With respect to the TOST’s size, given *α, θ, σ*_*ν*_, *ν* and *c*, the probability of declaring equivalence can be expressed as follows ^37^:

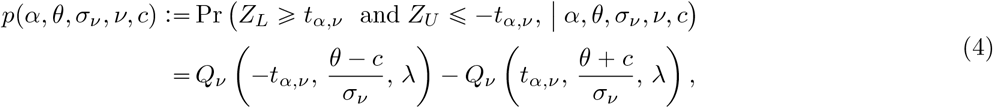

where 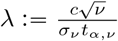 and *Q*_*ν*_ (*t, y, z*) corresponds to a special case of Owen’s Q-function ^38^ defined as

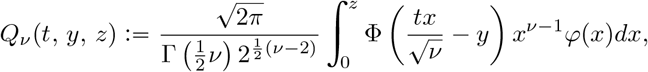

where *φ*(*x*) and Φ(*x*) denote the probability and cumulative distribution functions of a standard normal distribution, respectively.

Then, for given values of *α, σ*_*ν*_ and *ν*, the TOST’s size is defined as the supremum of (4), ^39^ and is given by

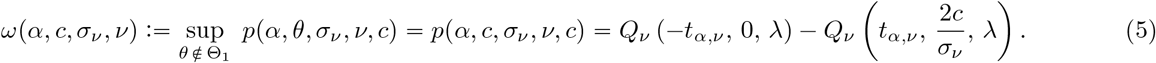

We can then deduce that the TOST is level-*α*, by noting that, for *σ*_*ν*_ >0, we have

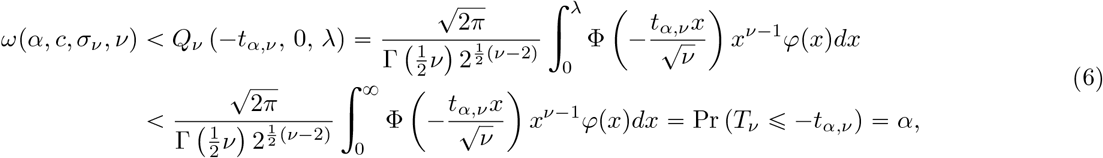

where *T*_*ν*_ denotes a random variable following a *t*-distribution with *ν* degrees of freedom, so that

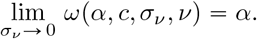

Thus, while the TOST is indeed level-*α*, it actually never achieves a size of *α*, except in the theoretical case of *σ*_*ν*_ = 0, as already highlighted by several authors. ^36^ When 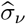 is small, the difference between the size and *α* is marginal, but as 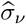 approaches or exceeds *c*/*t*_*α,ν*_, this difference increases, leading to a high probability of the TOST failing to detect equivalence when it exists. As a solution to this issue, we suggest an alternative approach, the *α*-TOST, that corrects the size of the TOST for a large range of values of 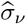and still allows to assess equivalence by means of confidence intervals, as depicted in Figure 5 of Section 4.

### 2.2 The *α*-TOST

A corrected version of the TOST can theoretically be constructed by adjusting the significance level and using *α*^*^ instead of *α* in the standard TOST procedure, where

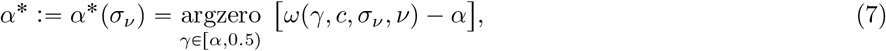

with *ω*(*γ, c, σ*_*ν*_, *ν*) defined in (5). The dependence of *α*^*^ on *α* and *ν* is omitted from the notation as these quantities are known. A similar type of correction was also used to amend the significance level of the SABE procedure by Labes and Schütz ^28^ and Ocaña and Muñoz ^35^ (see also Palmes et al. ^40^ for power adjustment). However, in these cases, the corrected significance level was reduced (instead of increased like in (7)) so that the size does not exceed the significance level of *α*. The aim of these corrections is therefore not the same as the one proposed here. Furthermore, the size of the *α*-TOST is guaranteed to be exactly *α* when *σ*_*ν*_ is known, which is not the case for these competing methods.

In Appendix A, we demonstrate that the existence of *α*^*^ relies on a simple condition that is satisfied in most settings of practical importance. In particular, this requirement can be translated into a maximal value for the estimated standard error 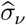, that is 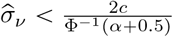. Moreover, since, *α*^*^(*σ*_*ν*_ ) is a population size quantity as it depends on the unknown quantity *σ*_*ν*_, a natural estimator for its sample value is given by

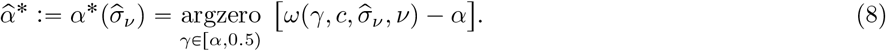

Hence, in practice, based on the (estimated) corrected significance level 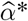, the *α*-TOST procedure rejects the non-equivalence null hypothesis in favour of the equivalence one at the significance level *α*, if 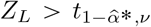 and 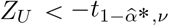. In Appendix B, we study the asymptotic properties of *α*^*^ and show that 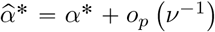. Informally, this result implies that the uncertainty associated to *α*^*^ is (asymptotically) negligible compared to the uncertainty associated to 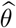 and 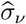as these terms have slower convergence rates in that 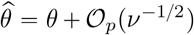 and 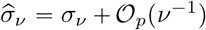. This result also suggests that the *α*-TOST procedures based on *α*^*^ or on 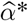are expected to provide very similar finite sample performances.

In Section 3, we consider an extensive Monte Carlo simulation study to compare the empirical performances of different methods when *σ*_*ν*_ needs to be estimated. For the 10^4^ simulation settings we considered (i.e., 100 values for *σ*_*ν*_ and 100 for *ν* covering most combinations of interest, see Simulation 2 in Table 1), we find that the empirical size of the *α*-TOST is generally closer to the nominal level *α* in comparison to the other methods (see Figure 2 in Section 3). We also find that in less that 1% of the settings, the *α*-TOST procedure can be slightly liberal, with a maximal empirical size of 0.05311 (see Figure 9 in Appendix D). However, this behaviour can mostly be explained by the randomness associated to our large scale simulation.

**Table 1:** Parameter values used in each simulation, where *c* denotes the tolerance limit, *ν* the number of degrees of freedom, *θ* the target parameter and *σ*_*ν*_ its standard deviation, *α* the target significance level and *B* the number of Monte Carlo samples per simulation.

| Simulation |  |  |  |  |
| --- | --- | --- | --- | --- |
|  | 1 | 2 | 3 | 4 |
| $c$ | $\log(1.25) \approx 0.2231$ | | | |
| $\nu$ | 15, 30, 45 | 5, 6, 7, …, 100, 250, 500, 750, 1000 (100 values) | | 45 |
| $\sigma_\nu$ | 0.08, 0.12, 0.16 | 100 evenly spaced values between 0.01 and 0.3 | | 0.08, 0.12, 0.16 |
| $\theta$ | 30 evenly spaced values between 0 and 0.26 | $c$ | 0 | 30 evenly spaced values between 0 and 0.26 |
| $\alpha$ | 0.05 | | | |
| $B$ | $10^5$ | | | |
| Design | General (canonical form) |  |  | Paired |
| Methods | TOST, α-TOST and δ-TOST | | | TOST, α-TOST, δ-TOST, SABE and cSABE |

Appendix E

The corrected significance level 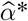 can easily be computed using the following iterative approach. At iteration *k*, with *k* ∈ ℕ, we define

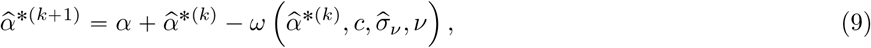

with *ω*(*α, c, σ*_*ν*_, *ν*) given in (5) and where the procedure is initialized at 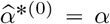. This simple iterative approach converges exponentially fast to 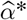 as it can be shown that

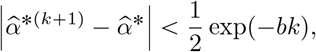

for some positive constant *b* (see Appendix C for more details).

Finally, since the conclusion of *α*-TOST considers an interval computed using a smaller value than *t*_*α,ν*_ compared to the TOST, the *α*-TOST rejection interval is necessarily larger than its TOST counterpart as 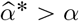. This implies that the *α*-TOST is uniformly more powerful than the TOST, and explains cases like the one encountered in the porcine skin case study presented in Section 4, in which equivalence is declared using the *α*-TOST but not with the TOST (which has an empirical power of zero given 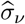 ).

## 3 Simulation Study

In this section, we conduct an extensive Monte Carlo simulation study with parameters settings per simulation reported in Table 1. Simulations 1 to 3, performed under the canonical form defined in (1) and therefore valid in a wide range of settings, assess the empirical performances of the *α*-TOST and compare them to the ones of the standard TOST and *δ*-TOST methods, where the latter, defined below, considers a correction on the equivalence limits rather than on the level to reach a size of *α*. Simulation 4, presented in Appendix E, investigates the empirical performances of these methods with the ones of the design-specific SABE and corrected SABE, where the latter consists in an adjustment on both the level and the equivalence limits. In that simulation, we consider a paired design setting that is closely related to the example considered in our case study and that allows us to estimate the within-subject variability of the reference treatment required by SABE-like methods. All simulations consider a target significance level of 5%, a value of *c* equal to logp1.25q and 10^5^ Monte Carlo samples per configuration.

Formally, the *δ*-TOST is defined as follows

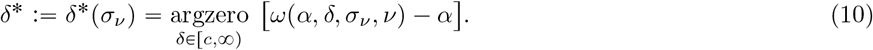

Using the same arguments as in Appendix A, we can easily demonstrate that a unique solution always exists, regardless of the value of the standard error for the *δ*-TOST. However, an exponentially fast iterative algorithm cannot be used to find the solution for this method. This highlights an important practical advantage of the *α*-TOST over the *δ*TOST and, to a larger extent, over the corrected SABE, which relies on both Monte Carlo integration and numerical optimisation procedures to define its correction (see Appendix E).

In our simulations, the empirical performances of the TOST, *α*-TOST and *δ*-TOST are defined using the following steps:

### 1. Simulation

for a given Monte Carlo sample *b* = 1, …, *B*:

a. simulate a value for 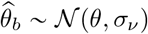 given the values *θ* and *σ*_*ν*_ of interest,
b. simulate a value for 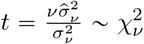 and set 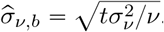.

### 2. Finite sample adjustments

for a given Monte Carlo iteration *b* = 1, …, *B*:

a. *α*-TOST:
  i. compute 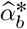 using (the algorithm associated to) (7) with 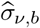,
  ii. compute 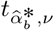 .
b. *δ*-TOST: compute 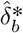 using (10) with 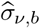.

### 3. Empirical probability of declaring equivalence

a. TOST:

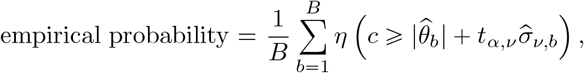

where *η*(·) denote the indicator function with *η*(*A*) = 1 if *A* is true and zero otherwise,
b. *α*-TOST: same as the TOST in Step 3(a) but replacing *t*_*α,ν*_ by 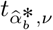
c. *δ*-TOST: same as the TOST in Step 3(a) but replacing *c* by 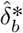.

Simulation 1 investigates the probability of declaring equivalence for varying values of *θ* allowing to study both the power and the size of each methods for combinations of selected values of *ν* and *σ*_*ν*_. Simulation results are presented in Figure 1, which shows, for each method of interest, the empirical probability of declaring equivalence as a function of *θ* for different combinations of values of *ν* (rows) and *σ*_*ν*_ (columns). For small values of *σ*_*ν*_, the empirical performance of all methods are similar. For moderate to large values of *σ*_*ν*_, we can note that the TOST is conservative, with an empirical size far smaller than the nominal level *α* = 5% when *θ= c*, and that it quickly reaches an empirical power of 0 for large values of *σ*_*ν*_. On the other hand, the *α*-TOST and *δ*-TOST have a higher power throughout, are generally size-*α* but are a bit conservative for large values of *σ*_*ν*_ and relatively small values of *ν*. This deviation from the nominal level *α* for the *α*-TOST and *δ*-TOST is due to the estimation error induced by using 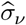 instead of *σ*_*ν*_ to construct an adjustment to the TOST. However, our simulation results confirm that such adjustment, either on the level or the equivalence bounds, considerably improves both the size and power in finite samples, especially with larger *σ*_*ν*_ where it prevents it from becoming 0. Moreover, this simulation suggests that the *α*-TOST outperforms the *δ*-TOST, indicating that an adjustment on the level provides both a more accurate and a more powerful test than an adjustment on the equivalence bounds.

**Figure 1:**
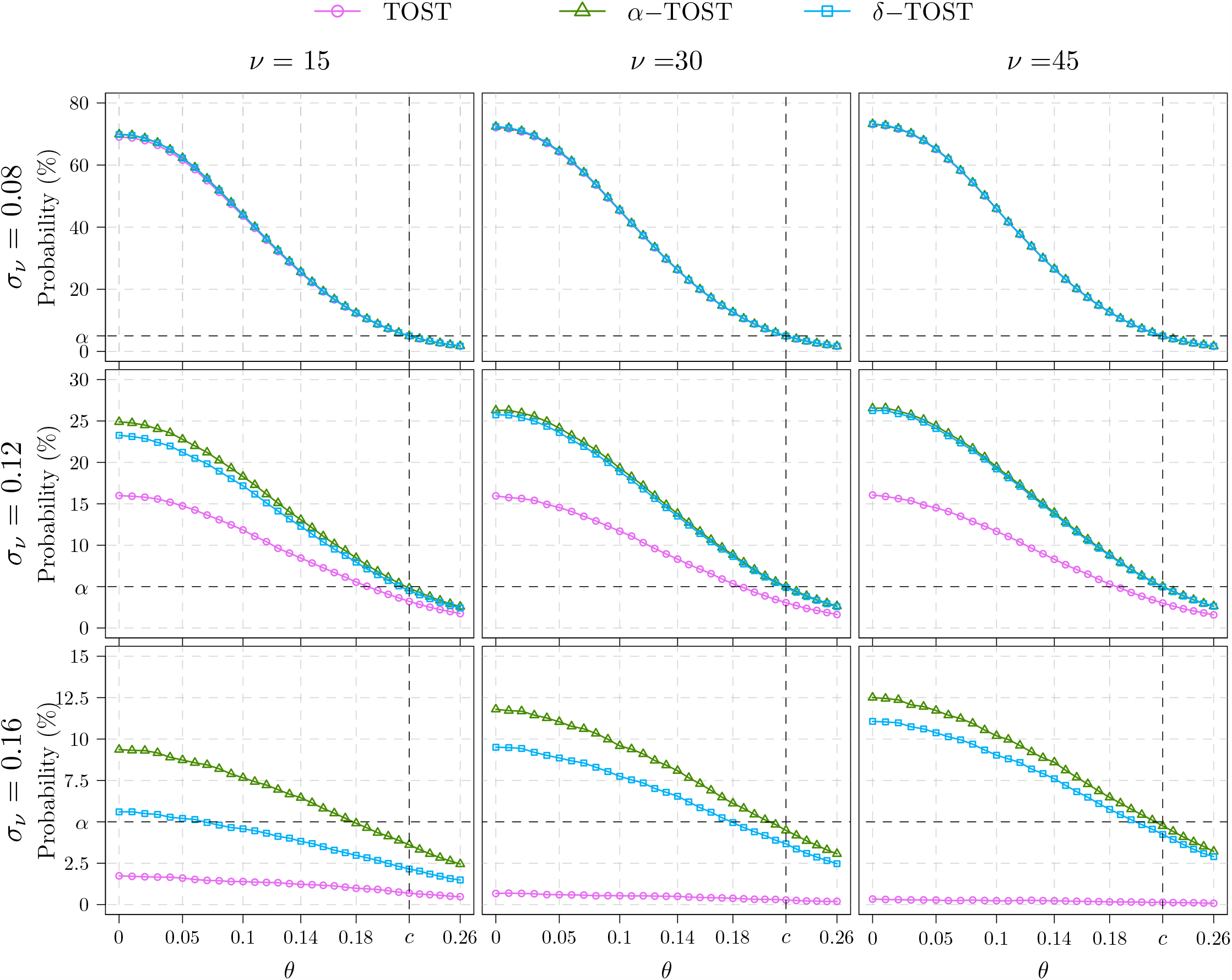
Empirical probability of declaring equivalence (*y*-axis) as a function of *θ* (*x*-axis), *ν* (columns) and *σ*_*ν*_ (rows), for the TOST (pink circles), the *α*-TOST (green triangles), and the *δ*-TOST (blue squares). Refer to the settings of Simulation 1 in Table 1 for details. In all configurations considered here, *α*-TOST shows a similar or greater power than the TOST and *δ*-TOST while remaining more accurate in terms of empirical size.

**Figure 2:**
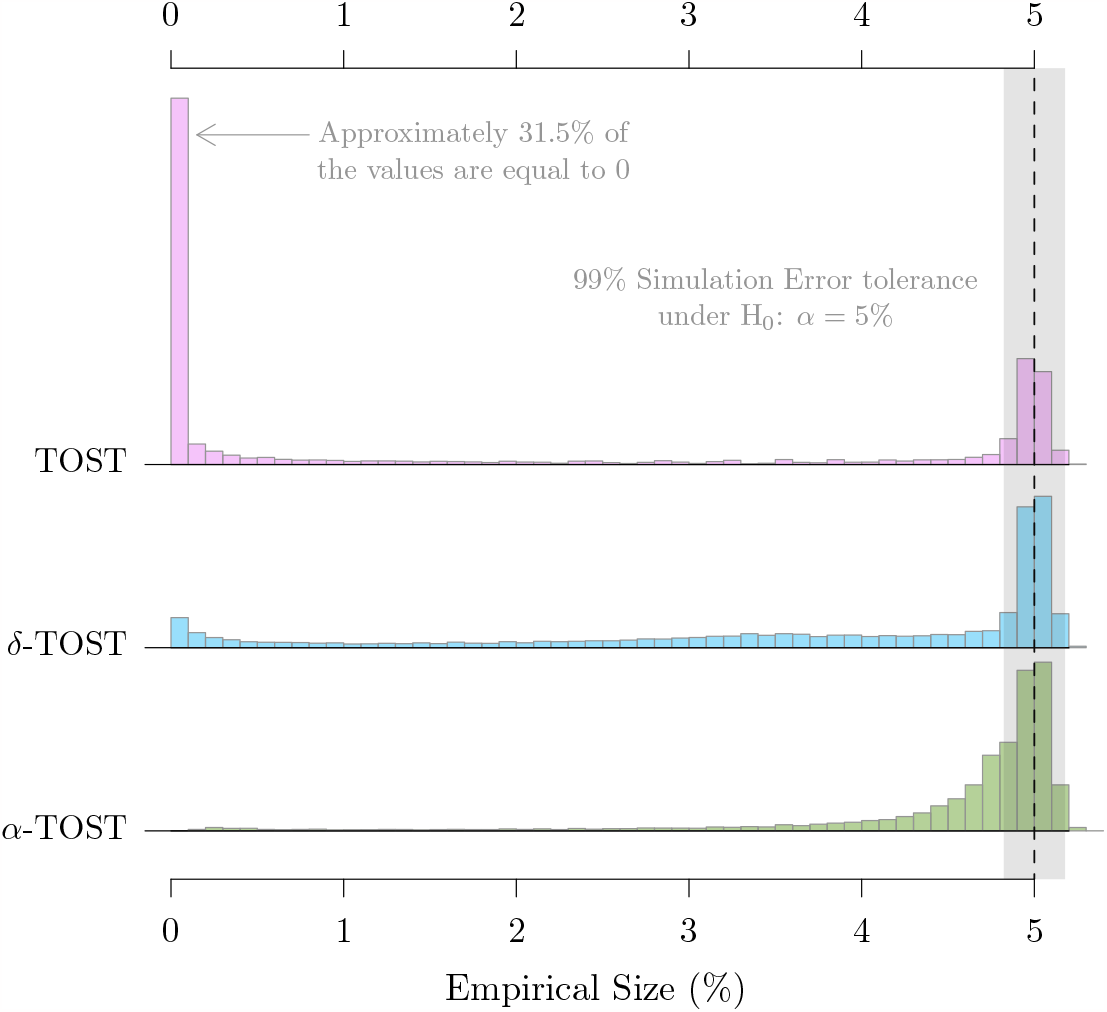
Histograms of the empirical size (%) of the TOST (first line), the *δ*-TOST (second line) and *α*-TOST (third line), computed from the results displayed in Figures 7, 8 and 9 in Appendix D, respectively. Overall, the *α*-TOST maintains a size of *α* for a larger proportion of parameters’ in comparison to the other methods.

**Figure 3:**
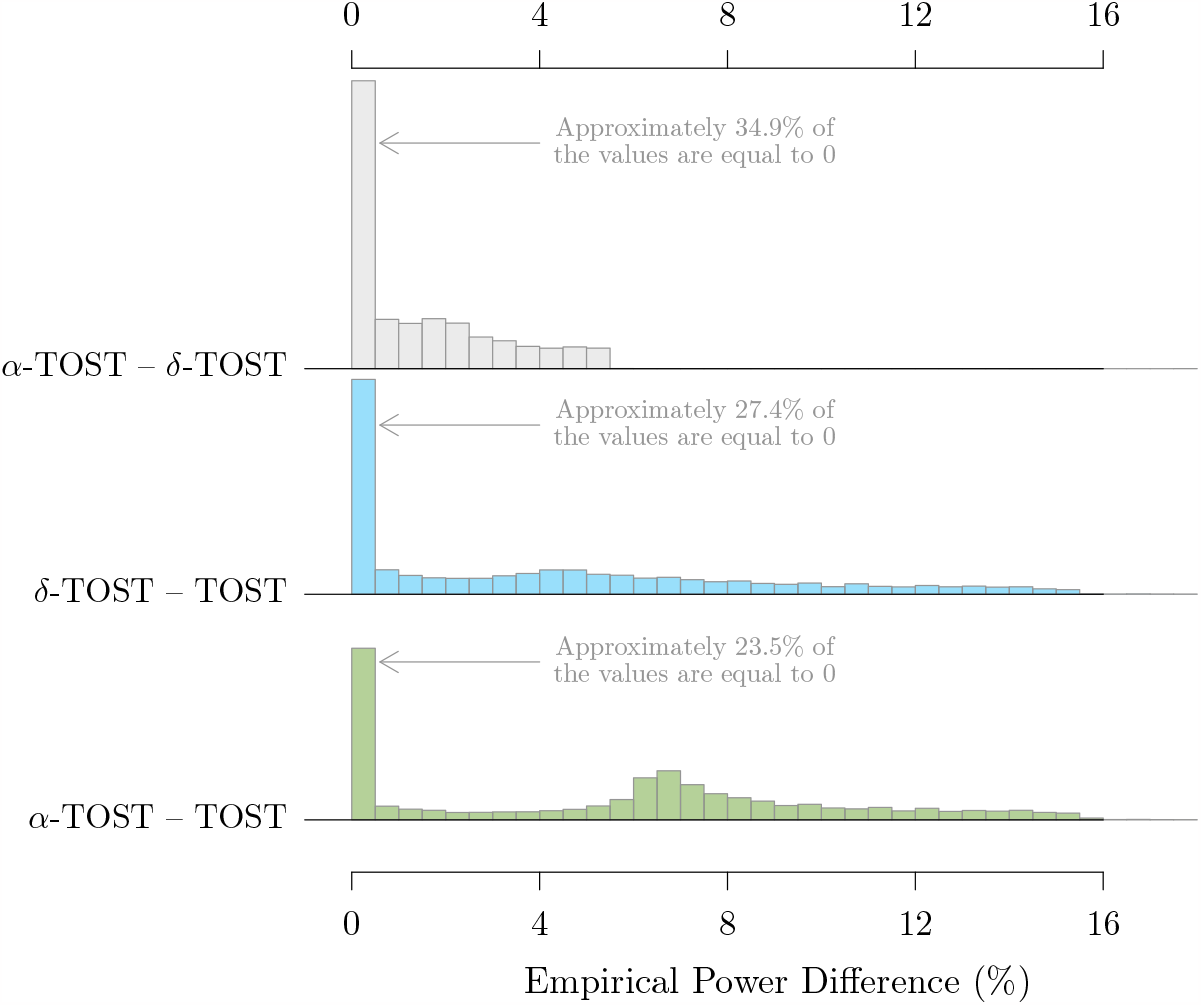
Histograms of empirical power differences (%) for each pair combinations of the TOST, the *δ*-TOST and *α*-TOST, computed from the results displayed in Figures 7, 8 and 9 in Appendix D. Overall, the *α*-TOST is the most powerful, followed by the *δ*-TOST then by the TOST.

**Figure 4:**
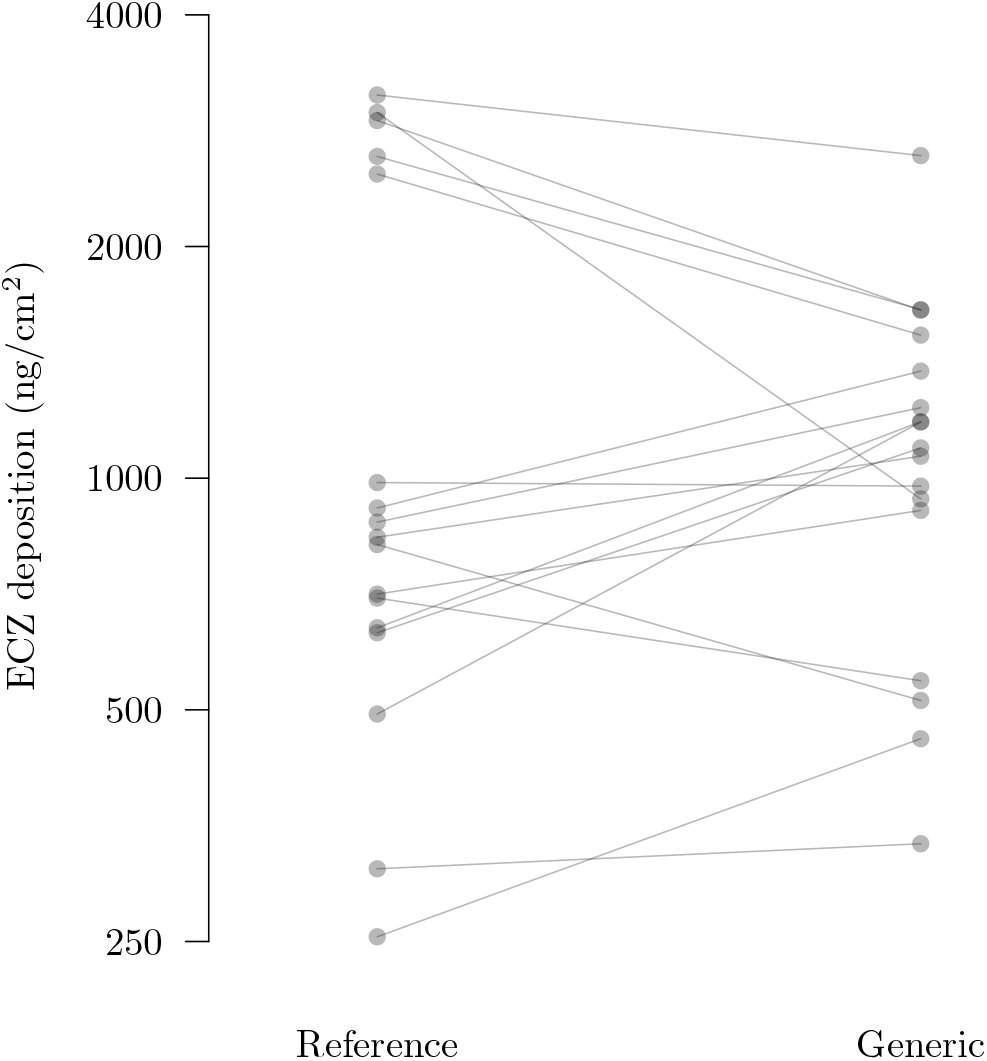
Econazole nitrate deposition levels (*y*-axis) measured using the reference and generic creams (*x*-axis) on 17 pairs of comparable porcine skin samples (lines).

**Figure 5:**
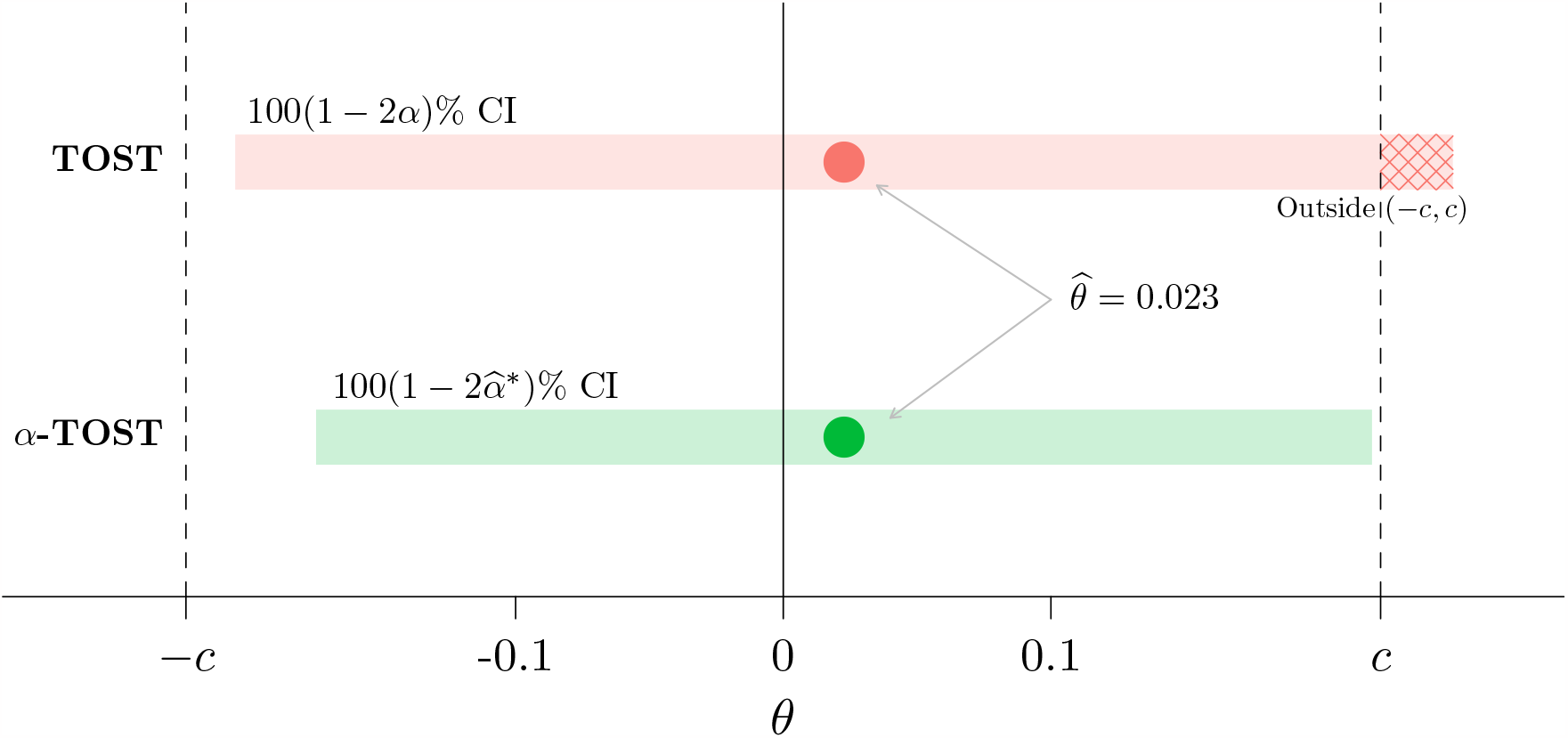
100 (1 −2^*α*^) % and 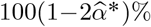 confidence intervals of the TOST and *α*-TOST procedures for the mean of the paired *log* differences in ECZ levels obtained with the reference and generic creams with *α* = 5% and 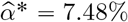. The dashed vertical lines correspond to the used lower and upper bioequivalence limits with *c* log 1.25. Comparison of the CI of each approach to the bioequivalence limits leads to the declaration of bioequivalence for the *α*-TOST procedure and not for the classic TOST approach due to its CI upper limit exceeding *c* (hatched area).

**Figure 6:**
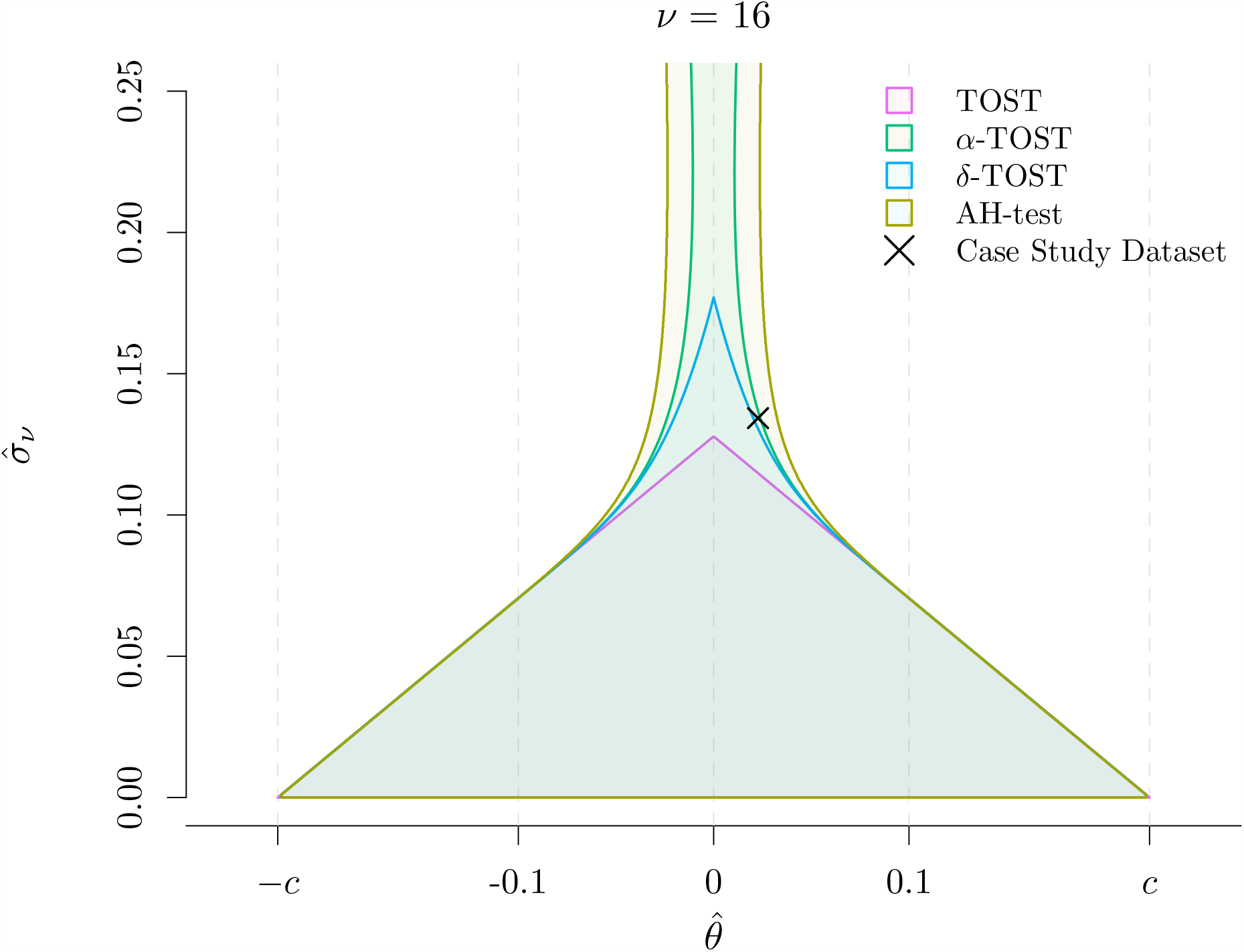
Bioequivalence test rejection regions as a function of 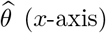 and 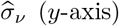 per method considered in Table 2 (coloured areas) showing combinations of values for 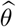 and 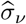 leading to equivalence declaration in the setting of the porcine skin dataset, i.e., with *c* =log(1.25) and *ν=* 16. The rejection regions of the different methods almost perfectly overlap for values of 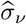 below 0.09 and differ for larger values. Regardless of 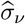, the TOST can’t declare bioequivalence for large values of 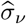(greater than0.12 here) and the *δ*-TOST for approximately 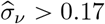, while the *α*-TOST and AH-test can, with the rejection region of the *α*-TOST embedded in the too liberal one of the AH-test. The symbol × represents the analysed data set in the acceptance/rejection regions where 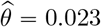 and 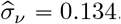.

Simulations 2 and 3, respectively performed with *θ=* log(*c*) and *θ* =0, investigate the empirical size and power for 10^4^ settings defined as combinations of 100 values of *ν* and 100 values of *σ*_*ν*_ chosen to cover *most* cases of practical interest. Figures 7, 8 and 9 in Appendix D respectively show the results of Simulation 2 for the standard TOST, *α*-TOST and *δ*-TOST. Each figure consists in a heatmap displaying the empirical size, computed by replacing *σ*_*ν*_ and *θ* by realizations of their random variables in (1) to reproduce the parameter estimation process, for all combinations of values of *ν* and *σ*_*ν*_ of interest. Figure 7 shows that the TOST is size-*α* only for relatively small values of *σ*_*ν*_ (below 0.09), and that its size decreases abruptly as *σ*_*ν*_ increases to reach 0. We can note that the value of *ν* does not seem to have an important effect on the size of the TOST. In comparison, Figures 8 and 9 show that both the *δ*-TOST and *α*-TOST are size-*α* for a larger number of combinations of values for *σ*_*ν*_ and *ν* and that the probability of being size-*α* increases with *ν* for a given value of *σ*_*ν*_. A comparison of Figures 8 and 9 shows that the *α*-TOST is both more powerful and more accurate than the *δ*-TOST overall, a conclusion in agreement with results of Simulation 1. A look at the proportion of configurations with an empirical size significantly greater than *α* – as assessed by a two-sided binomial exact test at the 1% level performed on the results obtained on the 10^5^ Monte Carlo samples per setting – shows that the *α*-TOST procedure is slightly liberal in 0.9% of the configurations considered in Simulation 2, with a maximal empirical size of 0.05311, compared to 0.0528 for the TOST and *δ*-TOST. This behavior can largely be attributed to the randomness inherent to our large-scale simulation.

**Figure 7:**
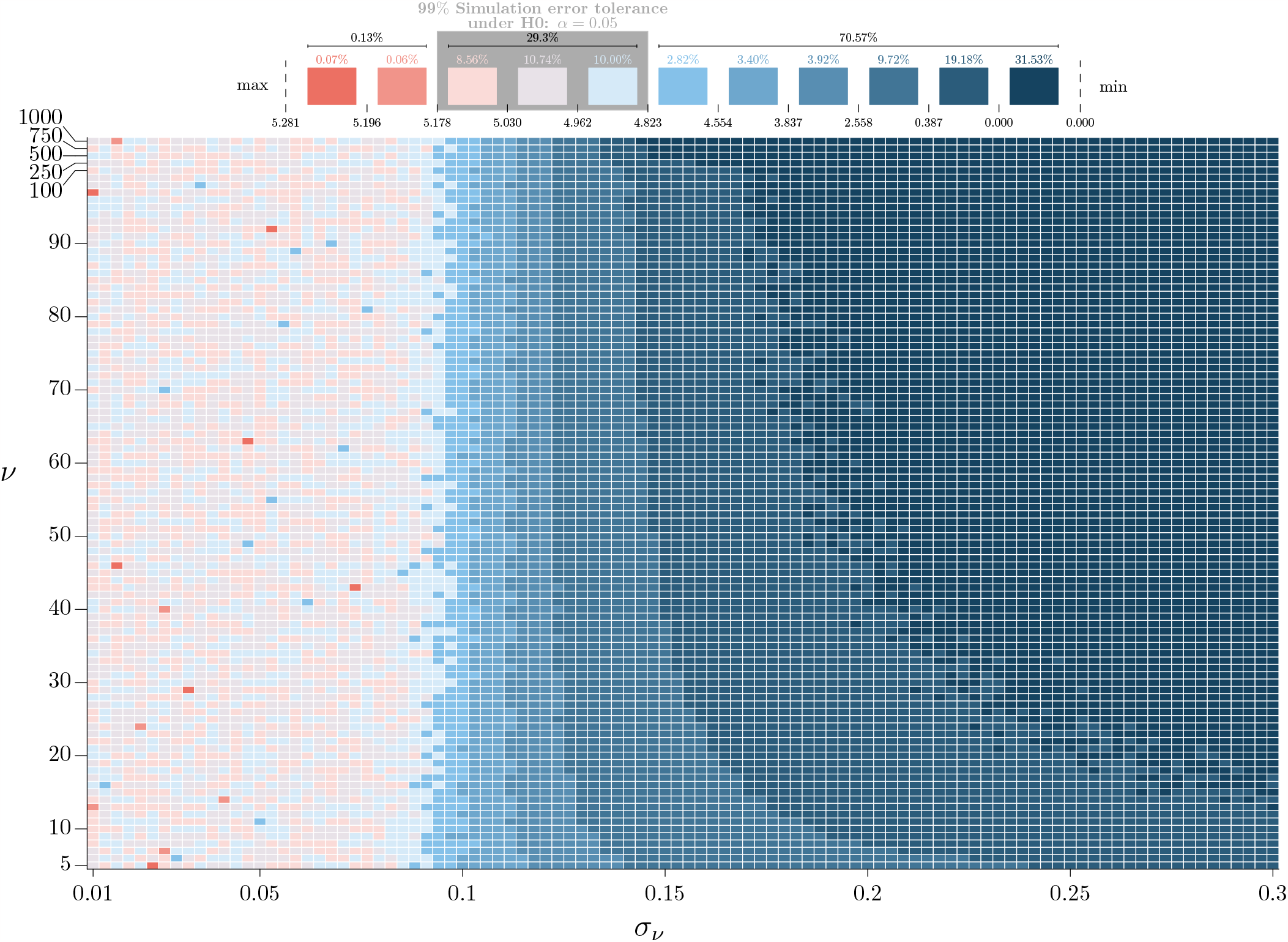
Heatmap representing the empirical size in % (color gradient) for the TOST, computed using the setting of Simulation 2 in Table 1, as a function of *σ*_*ν*_ (*x*-axis) and *ν* (*y*-axis). The lighter colors highlighted in the top legend correspond to the *α=* 5% nominal level, up to a simulation error assessed by a two-sided binomial exact test at the 1% level performed on the results obtained on the 10^5^ Monte Carlo samples per setting

**Figure 8:**
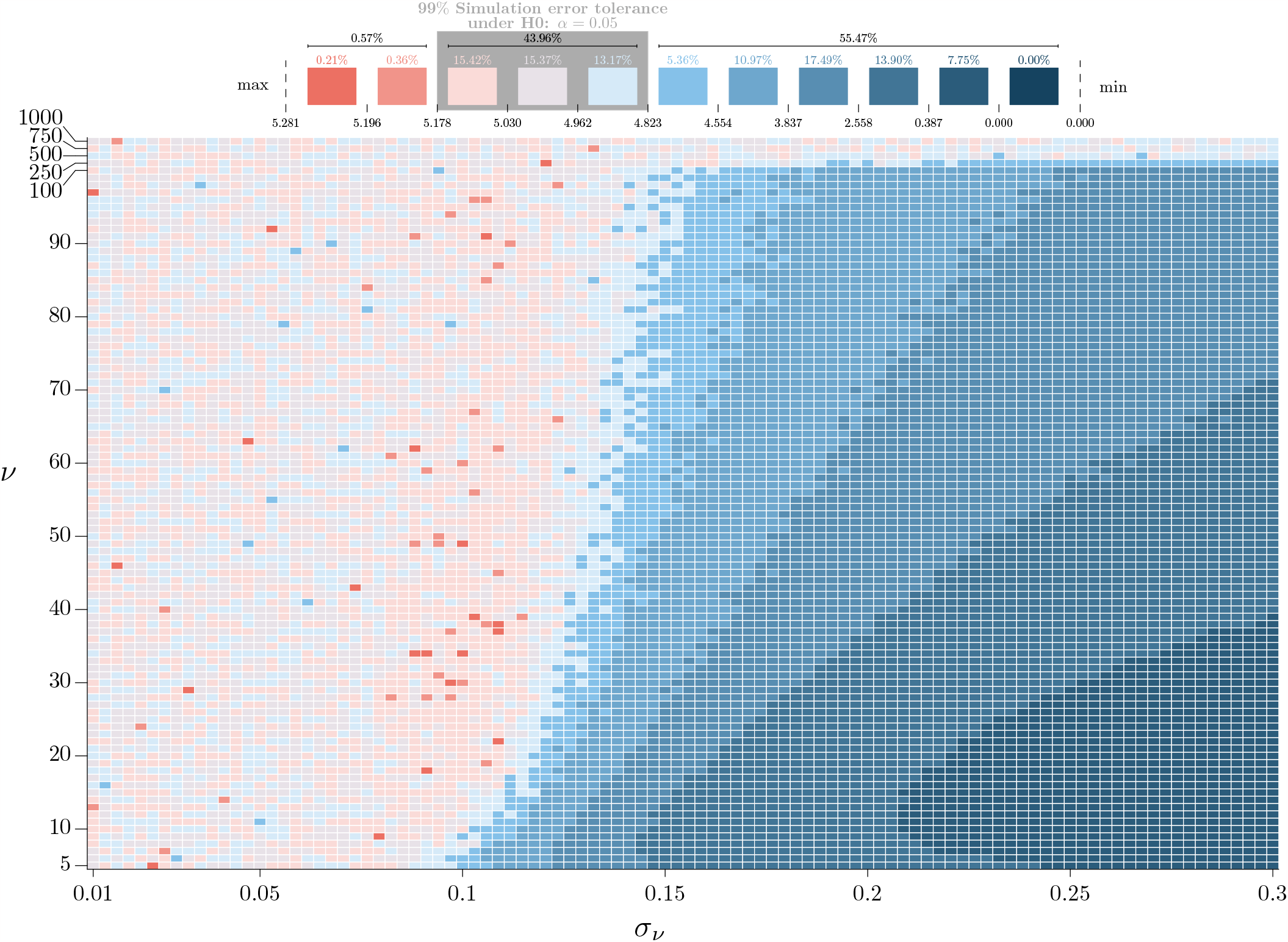
Heatmap representing the empirical size in % (color gradient) for the *δ*-TOST, computed using the setting of Simulation 2 in Table 1, as a function of *σ*_*ν*_ (*x*-axis) and *ν* (*y*-axis). The lighter colors highlighted in the top legend correspond to the *α =*5% nominal level, up to a simulation error assessed by a two-sided binomial exact test at the 1% level performed on the results obtained on the 10^5^ Monte Carlo samples per setting

**Figure 9:**
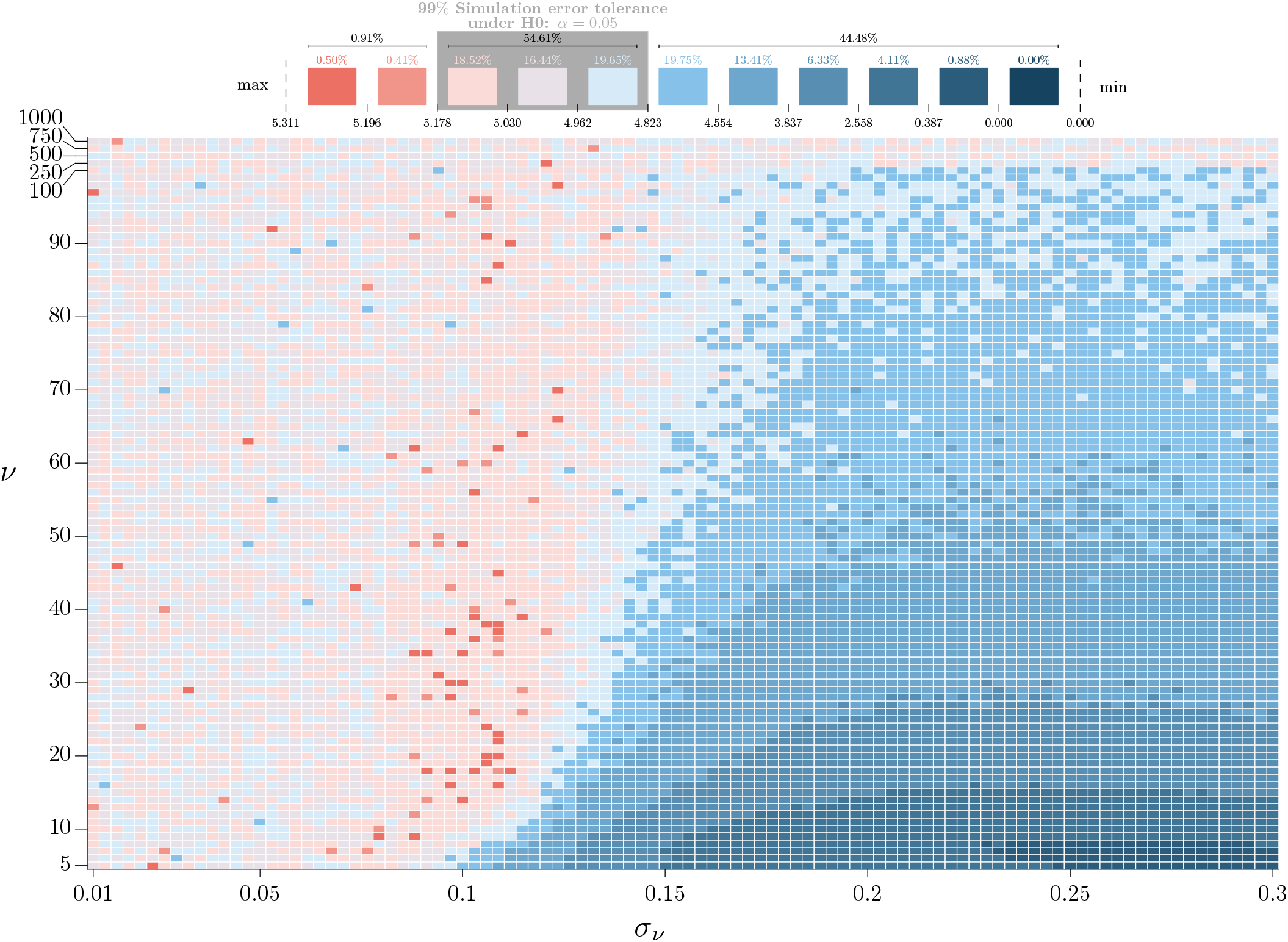
Heatmap representing the empirical size in % (color gradient) for the *α*-TOST, computed using the setting of Simulation 2 in Table 1, as a function of *σ*_*ν*_ (*x*-axis) and *ν* (*y*-axis). The lighter colors highlighted in the top legend correspond to the *α =*5% nominal level, up to a simulation error assessed by a two-sided binomial exact test at the 1% level performed on the results obtained on the 10^5^ Monte Carlo samples per setting.

Figure 2 summarises the results of Simulation 2 by displaying, for each method of interest, a histogram of the empirical sizes obtained over the 10^4^ configurations considered in the simulation. A comparison of these histograms clearly shows that the *α*-TOST has an empirical size closer to the nominal level *α* for a larger number of settings compared to the *δ*-TOST and standard TOST, which shows a large clump-at-zero (31.5%) corresponding to configurations with a power of 0.

The heatmaps in Figures 10, 11 and 12 in Appendix D show the results of Simulation 3 by displaying the power of each method for the same configurations as considered in Simulation 2. The results show that, over our 10^4^ configurations of interest, the *α*-TOST method is the most powerful, followed by the *δ*-TOST method and then the standard TOST. As expected, the power of all methods goes to one asymptotically as *σ*_*ν*_ → 0 (see Appendix B for details). Figure 3 summarises the results of Simulation 3 by displaying, for each pair of methods, the histogram of their differences in power for all configurations. The results show that the *α*-TOST is overall the most powerful, followed by the *δ*-TOST then by the TOST.

**Figure 10:**
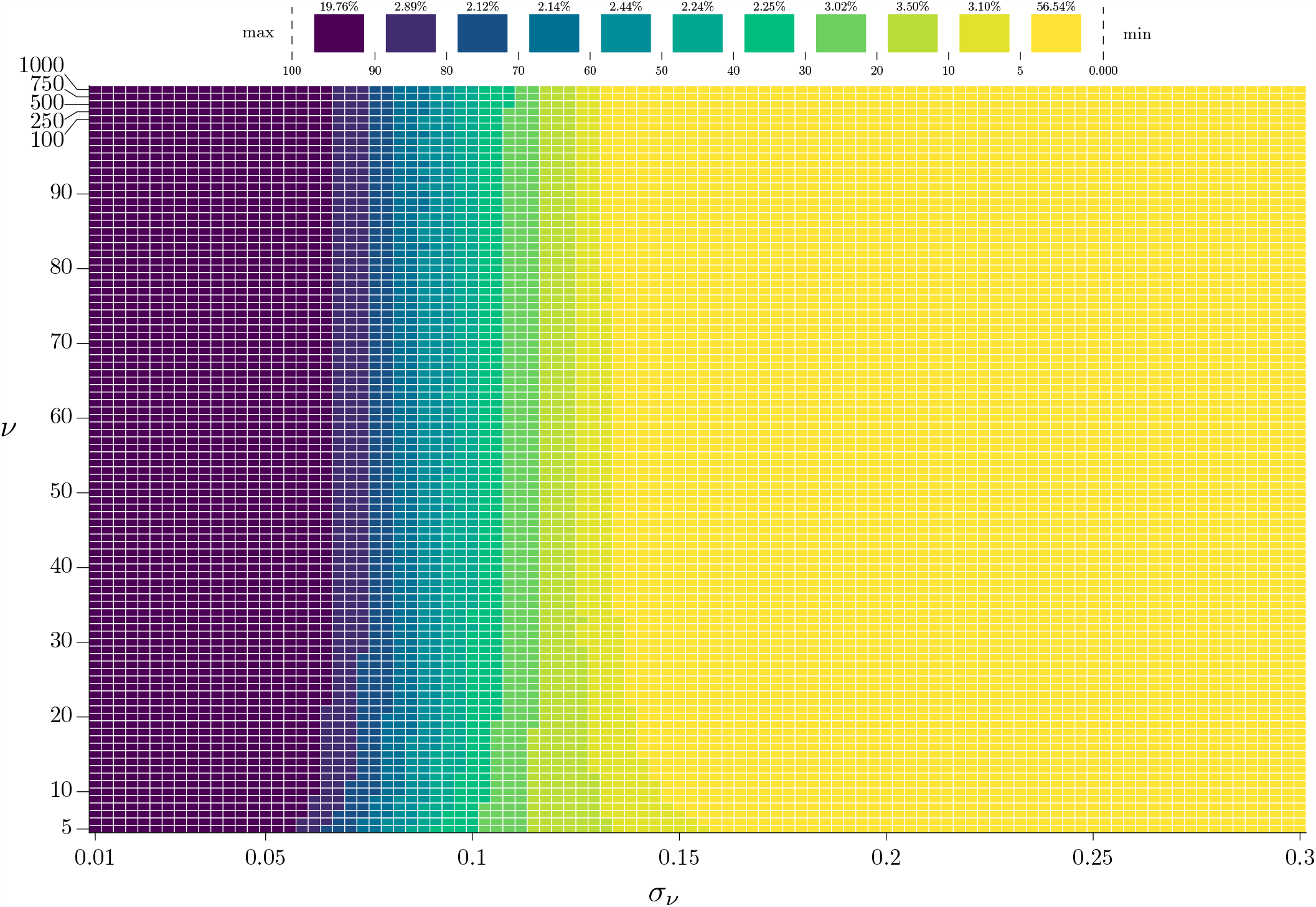
Heatmap representing the empirical power in % (color gradient) for the TOST, computed using the setting of Simulation 3 in Table 1, as a function of *σ*_*ν*_ (*x*-axis) and *ν* (*y*-axis).

**Figure 11:**
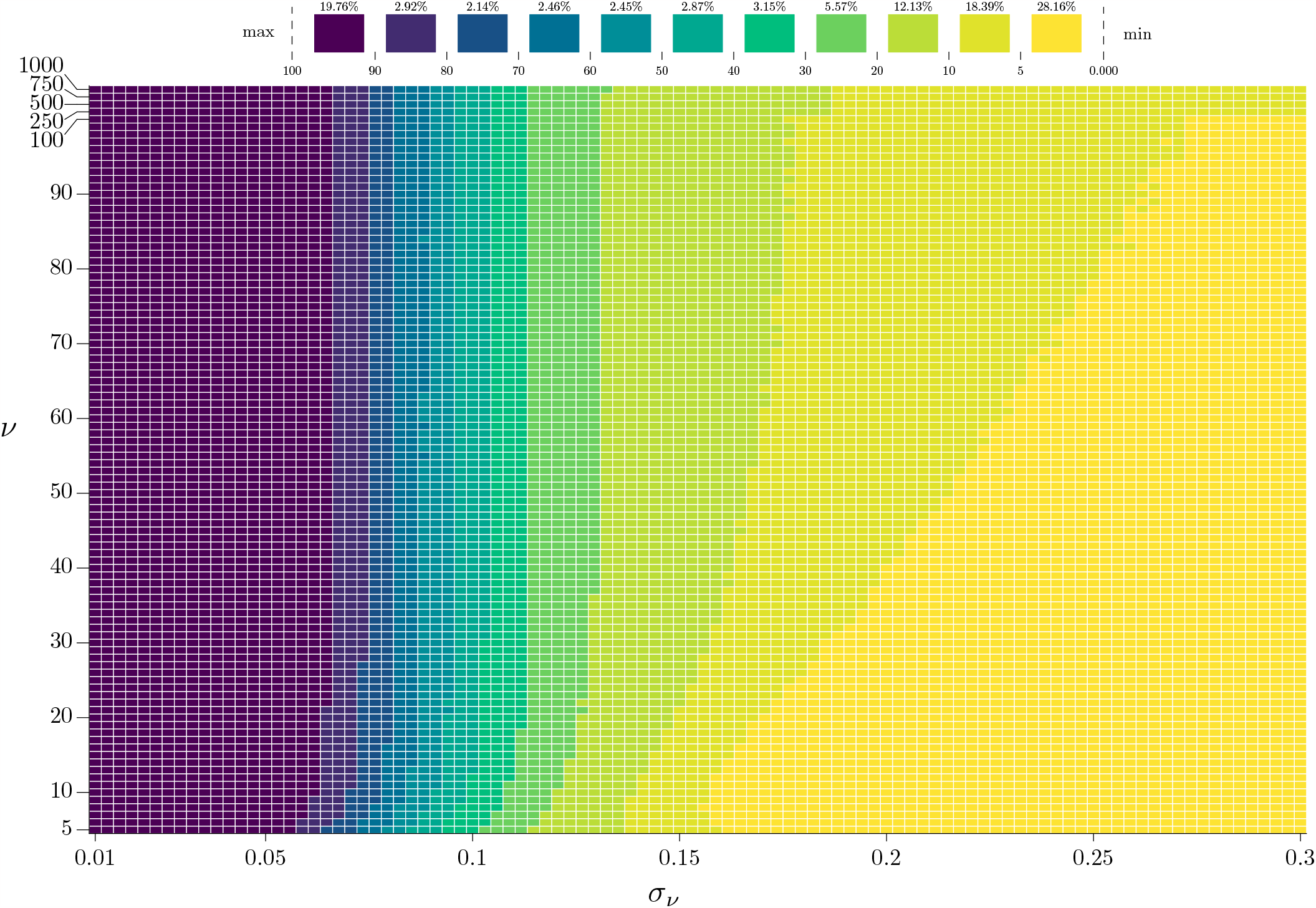
Heatmap representing the empirical power in % (color gradient) for the *δ*-TOST, computed using the setting of Simulation 3 in Table 1, as a function of *σ*_*ν*_ (*x*-axis) and *ν* (*y*-axis).

**Figure 12:**
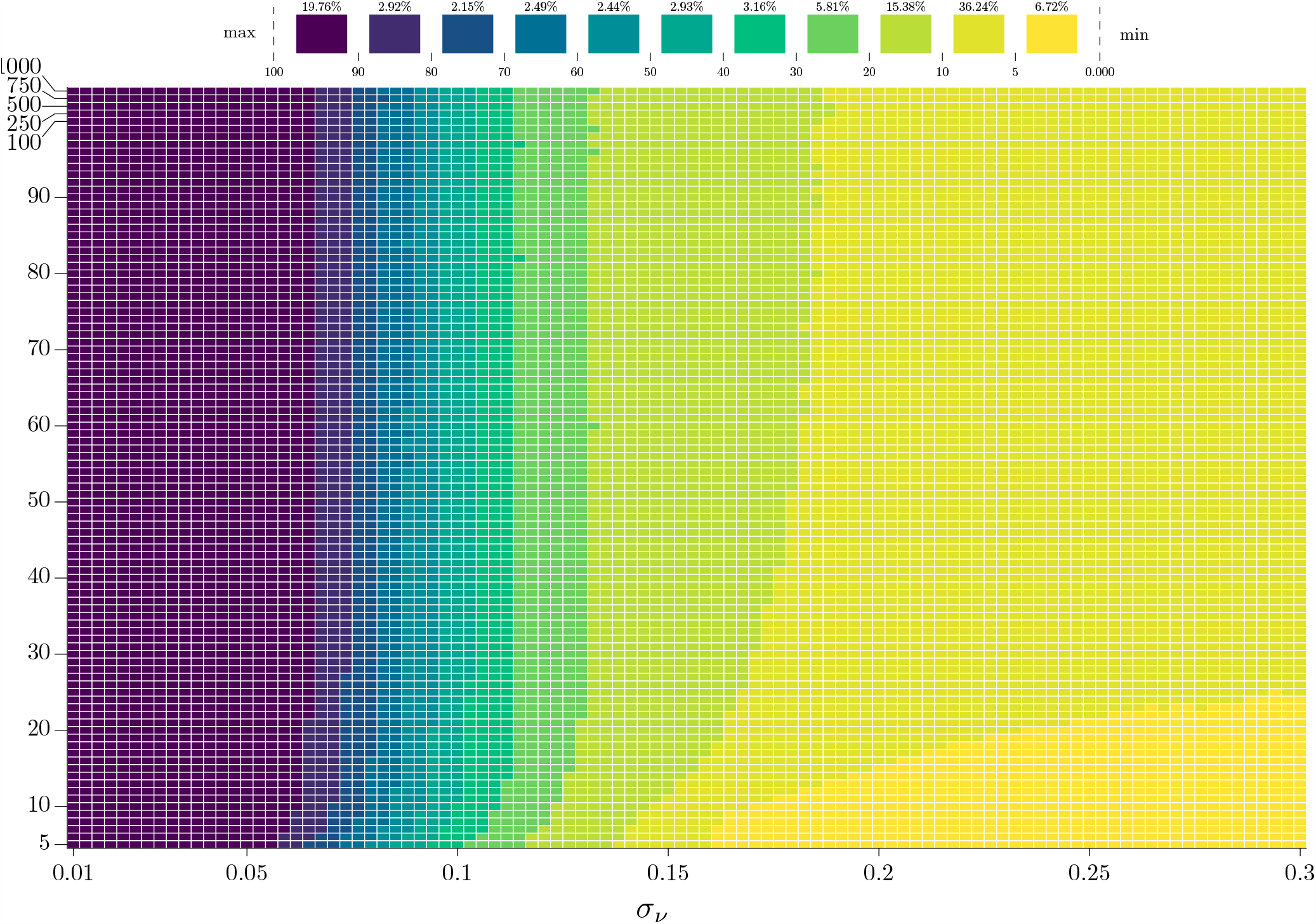
Heatmap representing the empirical power in % (color gradient) for the *α*-TOST, computed using the setting of Simulation 3 in Table 1, as a function of *σ*_*ν*_ (*x*-axis) and *ν* (*y*-axis).

In summary, the simulation studies considered here suggests that a correction of the TOST provides more power and better accuracy in finite samples, with considerably large improvements when *σ*_*ν*_ is large. Moreover, the *α*-TOST appears to provide a better performance than the *δ*-TOST, indicating that an adjustment on the level rather than on the equivalence bounds is preferable to enhance sample properties of equivalence tests. Results of Simulation 4, considering a paired study and additional correction methods, also suggest that adjusting the level of the TOST leads to better operating characteristics over competing methods, including the corrected SABE. More adequate and extensive simulations are needed though to compare these methods in the design-specific context required by regulatory agencies when assessing bioequivalence. Such simulations should consider the different adjustments proposed by regulatory agencies and are left for further research.

Finally, note that the idea of improving the size of the TOST is not new as Cao and Mathew ^41^ have proposed a correction based on the adjustment of the critical values defined as a non-increasing continuous function of the sample standard deviation to reduce the conservatism of the TOST. More particularly, they defined adjustment constants for specific values of 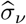 and used linear interpolation for the adjacent values. The lower panel of Figure 14 in Appendix F, compares the critical values obtained with the method of Cao and Mathew to the ones obtained by the *α*-TOST for different values of 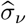and *ν*. We can note that, for all values of *ν* considered here and for values of 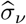 above 0.1, the corrected critical values ^41^ correspond to a piecewise version of the critical values obtained with the *α*-TOST when *ν* is large. Therefore, their correction appears to be an approximation of the *α*-TOST, evaluated asymptotically, i.e., at *ν* → ∞.

## 4 Evaluation of Bioequivalence for Econazole Nitrate Deposition in Porcine skin

Quartier et al. ^42^ studied the cutaneous bioequivalence of two topical cream products: a Reference Medicinal Product (RMP) and an approved generic containing econazole nitrate (ECZ), an antifungal medication used to treat skin infections. The evaluation of the putative bioequivalence is based on the determination of the cutaneous biodistribution profile of ECZ observed after application of the RMP and the generic product. The dataset we analyse in this section consists in 17 pairs of comparable porcine skin samples on which measurements of ECZ deposition were collected using both creams. Figure 4 presents the data, collected via a simple paired design, in which each pig delivered two skin samples respectively treated with one of the two drugs of interest. Such designs, possibly attractive for studies not involving regulators, are *stricto sensu* incompatible with the use of design-specific SABE-like corrections ^26^ and therefore interesting to showcase the advantages of our design-agnostic method. In order to assess bioequivalence of both topical treatments, the TOST and *α*-TOST procedures, based on a paired *t*-test statistic considering ECZ levels on the logarithmic scale, are conducted using *c* = *δ*_*U*_ = −*δ*_*L*_ = log(1.25) ≈ 0.223. Although the way to define bioequivalence limits for topical products is still being discussed, ^43^ we believe the chosen limits to be reasonable. ^42^

Figure 5 shows the CIs corresponding to both approaches. The 100(1 − 2^*α*^)% TOST confidence interval for the mean of the paired differences in ECZ levels equals [−0.204, 0.250], given that 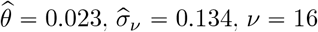 and *α* = 5%. As its upper bound exceeds the upper bioequivalence limit, the classical TOST procedure does not allow us to conclude that the topical products are (on average) equivalent. To reach a size of 5%, the *α*-TOST procedure uses in this case a significance level of 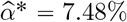 leading to a confidence interval of [−0.166, 0.211]. This CI being strictly embedded within the (−*c, c*) bioequivalence limits, the *α*-TOST procedure allows to declare bioequivalence, hence illustrating the increase in power induced by the increased significance level considered to reach a size of 5%. Note that in this case, given 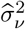and *ν*, the empirical power of the TOST is zero (regardless of *θ*) as *t*_0.05,16_ *σ*_*ν*_ > *c*, where *t*_*α,ν*_ denotes the upper quantile *α* of a *t*-distribution with *ν* degrees of freedom; see Appendix E. Since the *α*-TOST guarantees a size of *α* (for all sample sizes), the conclusion brought in by the *α*-TOST is more trustworthy.

To gain additional insight into the benefits conferred by our approach, we also compare the characteristics and conclusion of the *α*-TOST to other available methods in Table 2 as well as their rejection region as a function of 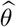 and 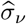 in Figure 6. We considered here the AH-test, the TOST, *α*-TOST and *δ*-TOST. The AH-test does not satisfy the IIP, but represents a good proxy for the other tests without this property and is relatively easy to implement. Among the level-*α* tests, the *α*-TOST is the only one leading to bioequivalence declaration.

**Table 2:** Bioequivalence declaration (yes/no) for the econazole nitrate deposition in porcine skin data using the AH-test, TOST, *α*-TOST and *δ*-TOST. The estimated parameter values are 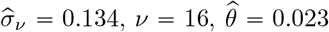 and *α =*5%. The columns IIP, Level-*α* and Size-*α*, respectively indicate if each method satisfies the IIP, if its size is bounded by *α* and if its size is exactly *α*. The symbol ^*^ specifies that the property is valid when the standard error *σ*_*ν*_ is known.

| Method | IIP | Level-α | Size-α | Bioequivalence Declaration |
| --- | --- | --- | --- | --- |
| AH-test | no | no | no | Yes |
| TOST | yes | yes | no | No |
| α-TOST | yes | yes* | yes* | Yes |
| δ-TOST | yes | yes* | yes* | No |

Figure 6 shows the combinations of values for 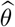 and 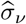 leading to bioequivalence declaration in the setting of the porcine skin dataset, i.e., with *c* = log(1.25) and *ν* =16. The rejection regions of the different methods almost perfectly overlap for values of 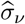 below 0.09 and differ for larger values. Regardless of 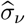, the TOST cannot declare bioequivalence for large values of 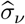 (greater than approximately 0.12 here) and the *δ*-TOST for approximately 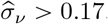, while the *α*-TOST and AH-test can, with the rejection region of the *α*-TOST embedded in the too liberal one of the AH-test. In Figure 6, the symbol × shows the values of 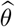 and 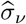 obtained in the porcine skin dataset. These coordinates lead to the declaration of (average) bioequivalence of the two topical products with the AH-test and *α*-TOST procedures, the later one only being size-*α*.

## 5 Discussion

The canonical framework treated in this paper is given in (1) and therefore concerns differences that can be assumed to be normally distributed (in finite samples), with a known finite sample distribution for 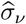. This framework covers a quite large spectrum of data settings, such as the standard two-period crossover experimental design, ^44^ and could be extended to include covariates to possibly reduce residual variance. Extensions to non-linear cases, such as for example binary outcomes, ^45,46,47,48,49^ would follow the same logic, but would require a specific treatment due to the nature of the responses and to the use of link functions. Such extensions also deserve some attention but are left for further research.

For sample size calculations, we could, in principle, proceed with the *α*-TOST, for given values of *c, θ* and *σ*_*ν*_. However, when considering high levels of power, the correction is negligible and we have *α*^*^ ≈ *α* as shown in Section 3, so that the sample size can be computed using the TOST, as implemented in standard packages. The *α*-TOST approach would then be used to assess equivalence and show its benefits when the observed value of *σ*_*ν*_ is unexpectedly large compared to the one considered in the sample size calculation either due to (lack of) chance or to an underestimated value obtained from a prior experiment.

## Acknowledgements

S. Guerrier is supported by the SNSF Grants #176843 and # 211007 as well as by the Innosuisse Grants #37308.1 IP-ENG and #53622.1 IP-ENG. Y. Boulaguiem and M.-P. Victoria-Feser are partially supported by the SNSF Grant #182684. D.-L. Couturier is partially supported by the Cancer Research UK grant C9545/A29580. Y. N. Kalia thanks the University of Geneva for teaching assistantships for J. Quartier and M. Lapteva, for providing financial support for the purchase of the Waters Xevo TQ-MS detector, and also thanks the Fondation Ernst and Lucie Schmidheiny and the Société Académique de Gené ve for providing equipment grants.

## Conflict of Interest

None declared.

## Data availability statement

The econazole nitrate deposition data as well as an implementation of the method proposed in this paper are available in the `cTOST` R package accessible on the GitHub repository: `yboulag/cTOST`.

## Appendix

### A. Existence of *α*^*^

In this section, we state the conditions for *α*^*^, defined in (7), to be a singleton. Fixing *α, c, σ*_*ν*_, and *ν*, we simplify our notation so that *ω* (*γ*) := *ω* (*γ, c, σ*_*ν*_, *ν*) and let

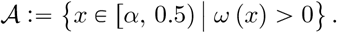

The function *ω*(*γ*), defined in (5), is continuously differentiable and strictly increasing in *γ* in *𝒜*. From (6), we have that *α* ⩾ *ω*(*α*). Thus, it is sufficient to show that 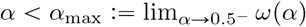, where *α* → 0.5^−^ denotes the limit from below 0.5, to ensure that *α*^*^ is a singleton. Let *T*_*ν,δ*_ denote a random variable following a non-central *t*-distribution with *ν* degrees of freedom and noncentrality parameter *δ* = 2*c*/*σ*_*ν*_. Then, we have

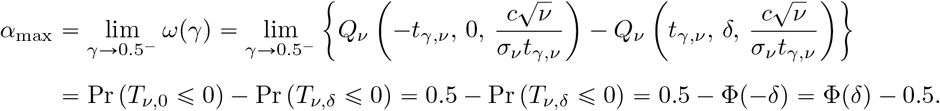

Thus, the condition *α* < *α*_max_ can be expressed as follows

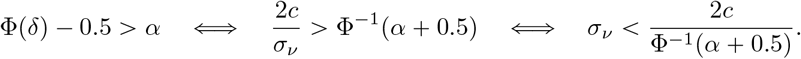

Therefore, the condition 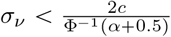 implies that *α* < *α*_max_ and consequently that *α*^*^ is a singleton. The existence of 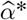 follows the same argument but replacing *σ*_*ν*_ by 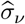 in our condition.

### B. Asymptotic Properties

In this section, we study the convergence rates of 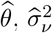 and 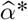. In particular, we show that

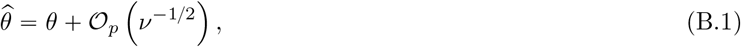

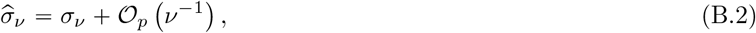

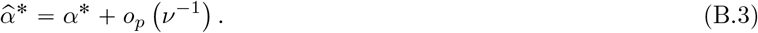

These results are based on the following standard regularity conditions. First, there exists a positive constant *σ*^2^ such that 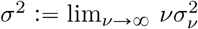. Second, the sequences

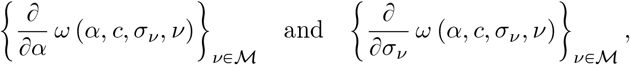

where *ℳ* ⊆ ℝ, converge uniformly in *α* and *σ*_*ν*_. This second condition implies by Theorem 7.17 of Rudin (1976) ^50^ that,

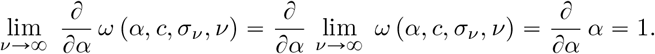

Similarly,

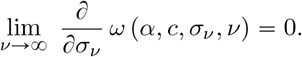

Using these regularity conditions, we start by proving (B.1). From (1), we have that 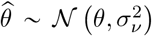 and, thus, by Markov’s inequality, for any *M* > 0, we obtain

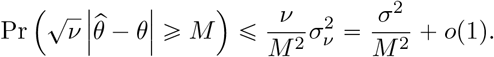

Therefore, we have 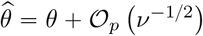, which verifies (B.1).

Next, we study the convergence rate of 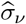. Let *Y* denote a random variable following a *χ* distribution with *ν* degrees of freedom, i.e., *Y* ∼ *χ*_*ν*_. We have

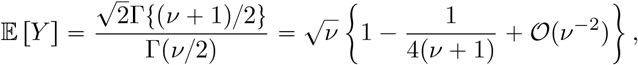

where Γ(·) is the gamma function, and where the second equality can be obtained using Stirling’s approximation for the Gamma function. Moreover, we have

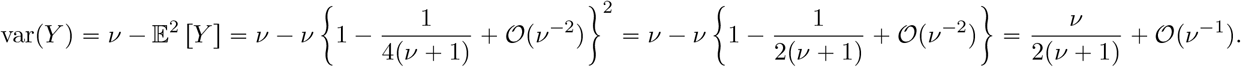

From (1), we have 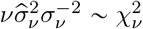 using Markov’s inequality, for any *M* > 0, we obtain

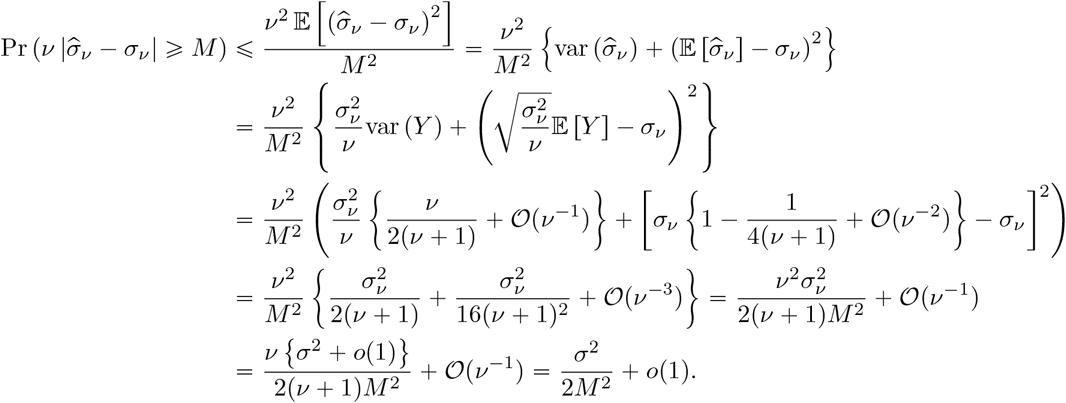

Thus, we have 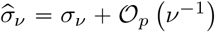, which verifies (B.2).

Finally, we consider the convergence rate of 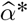. Using (B.2) and the continuity in *σ*_*ν*_ of *ω* (*α, c, σ*_*ν*_, *ν*), we have by the continuous mapping theorem that 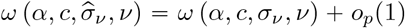, which by Lemma 5.10 of Van der Vaart (2000) ^51^ implies that

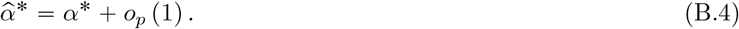

Next, there exists a point 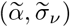 on the line segment from (α*, σ_*ν*_ to 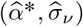 such that

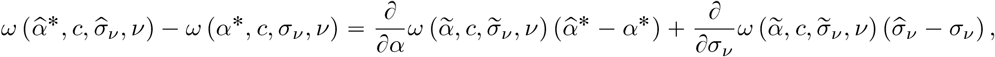

Where

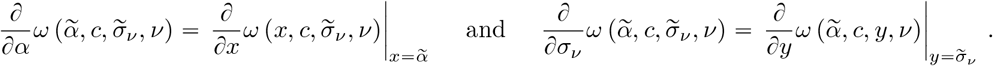

From (7) and (8), we have *ω* (*α*^*^, *c, σ*_*ν*_, *ν*) = *α* and 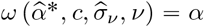, implying that

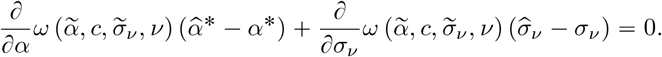

Since ^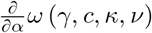^ converges uniformly, for all *ε* > 0, there exists *N*_*ν*_ > 0 such that for all *γ* ∈ [*α*, 0.5), for all *κ* ∈ ℝ _+_and for all *ν* ⩾ *N*_*ν*_, we have that 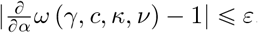. Thus, we have

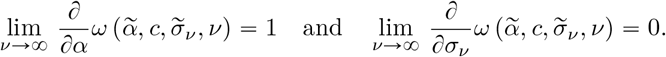

Consequently, we obtain

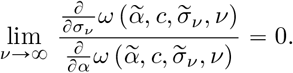

For sufficiently large *ν*, we have

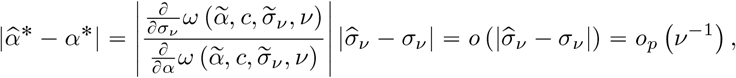

since, from (B.2), we have 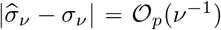. Therefore, we obtain 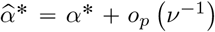, which verifies (B.3) and concludes the proof.

### C. Convergence Rate of the Iterative Approach for *α*^*^

Using the notation of Appendix A and for *γ* ∈ *𝒜*, we have that *ω* (*γ*) is continuously differentiable and such that 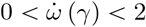, where

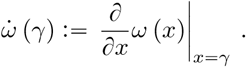

Next, we define

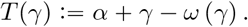

For all *α*_1_, *α*_2_ ∈ *𝒜*, we have the mean value theorem stating that

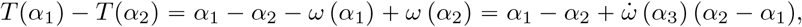

where *α*_3_ = *τα*_1_ +(1 − *τ* )*α*_2_ for some *τ* ∈ [0, 1]. Thus, we obtain

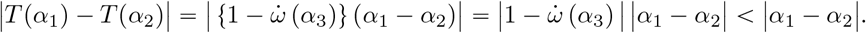

Then, using Kirszbraun theorem, ^52^ we can extend the function *T* (*γ*) with respect to *γ* ∈ *𝒜* to a contraction map from ℝ to ℝ. Thus, Banach fixed point theorem ensures that *T (α*^*(*k)*^)converges as *k* →8. We then define the limit of the sequence {*α*^*(*k*+1)^_*k* ∈ ℕ_ as *α*^*^, which is the unique fixed point of the function *T* (*γ*). Indeed, we have

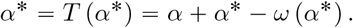

By rearranging terms, we have

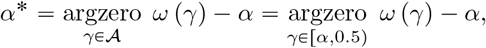

concluding the convergence of the sequence {*α*^*(*k*+1)^_*k* ∈ ℕ_ as *α*^*^. As a result, there exists some 0 < *ϵ* < 1 such that for *k* ∈ ℕ we have

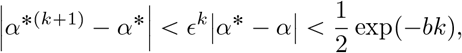

for some positive constant *b*.

### D. Empirical Size and Power Comparisons for the TOST, *α*-TOST and *δ*-TOST

In this section, we perform an extensive simulation study, for the evaluation of the empirical size and power of the *α*-TOST, compared to the TOST and *δ*-TOST, by varying the values of *ν* and *σ*_*ν*_ over a large grid. The power (i.e. when *θ* = 0) and the size (i.e. when *θ* = *c*) are computed by replacing both *σ*_*ν*_ and *θ* by realizations of their corresponding random variables in (1), i.e. reproducing the case of parameter estimation. The simulation settings we consider are given in Simulations 2 and 3 of Table 1 for the size and power respectively.

### E. Empirical comparisons with the corrected SABE

In this Section, we present a simulation study to compare the power and level of the TOST, *α*-TOST, *δ*-TOST, the EMA implementation of the SABE and its corrected version proposed by Labes and Schütz, ^28^ which they call the iteratively adjusted *α* of the Average BioEquivalence Expanding Limits (ABEL). The aim is to compare testing procedures that either correct for the level (*α*-TOST), for the equivalence limits (*δ*-TOST) or both (the corrected SABE) in a paired setting closely related to the case study presented in Section 4 and allowing to estimate the withinsubject variability of the reference treatment required by SABE-like methods. We should stress that the SABE and its corrected version are usually used with replicated cross-over designs ^26^ and that their use in a simple paired design can be viewed as a relaxation of the constraints imposed by regulatory authorities that still allows to validly investigate their finite sample properties. The corrected SABE method is implemented as defined by the EMA guidelines, by computing the size through Monte Carlo integration (10^5^ simulations), and applying a correction on the level to match the original one, *α*, only when the test is liberal given the adjusted (bio)equivalence limits. Note that, as the SABE procedure also requires the estimated 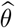 to lie inside the *original* equivalence bounds to declare bioequivalence, the size can only be computed using Monte Carlo simulations. The model we consider is given by

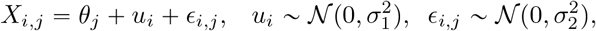

with *i* = 1, …, *n* and *j=* 1, 2. By taking the paired differences we obtain

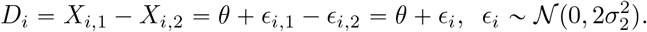

Thus, we have 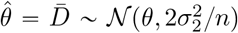. By defining 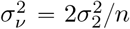 and *ν* = *n* − 1, we have 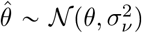 and ^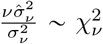.^ In this setting, the Coefficient of Variation (CV) is 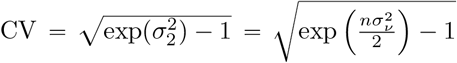, and we consider the following estimator

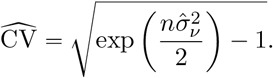

The parameters and settings considered in this Simulation are reported in Table 1 under Simulation 4. The first row of Figure 13 shows the empirical probabilities of declaring equivalence (y-axes) of each method (coloured lines) as a function of *θ* (x-axes) for two values *σ*_*ν*_ (columns) when *ν* = 45.Comparing the empirical size - obtained when *c* = *θ* - of the different methods, we can note that the TOST is quite conservative while the SABE is very liberal for the considered values of *σ*_*ν*_. The *α*-TOST, *δ*-TOST and cSABE are size-*α* when *σ*_*ν*_ = 0.12 with their respective empirical size lying inside the 99% simulation error tolerance interval of (4.84, 5.16), corresponding to *α=* 5% and *B* = 10^5^ Monte Carlo simulations. On the other hand, for a value of *σ*_*ν*_ = 0.16, none of these methods is size-*α* as all empirical sizes lie below the simulation error tolerance interval (with estimated values of approximately 0.0475, 0.0424 and 0.0357 for the *α*-TOST, the *δ*-TOST and the cSABE, respectively). The second row of the Figure 13 shows the difference in the probabilities of declaring equivalence of the *α*- and *δ*− TOST methods compared to the cSABE (y-axes) as a function of *θ* (x-axes) for two values of *σ*_*ν*_ (columns) when *ν* = 45. Empirically, we can note that, in terms of power, the *α*-TOST uniformly dominates the other two methods and the *δ*-TOST uniformly dominates the cSABE. This again suggests that an adjustment on the level of the TOST is the most effective way to improve the finite sample properties of equivalence testing.

**Figure 13:**
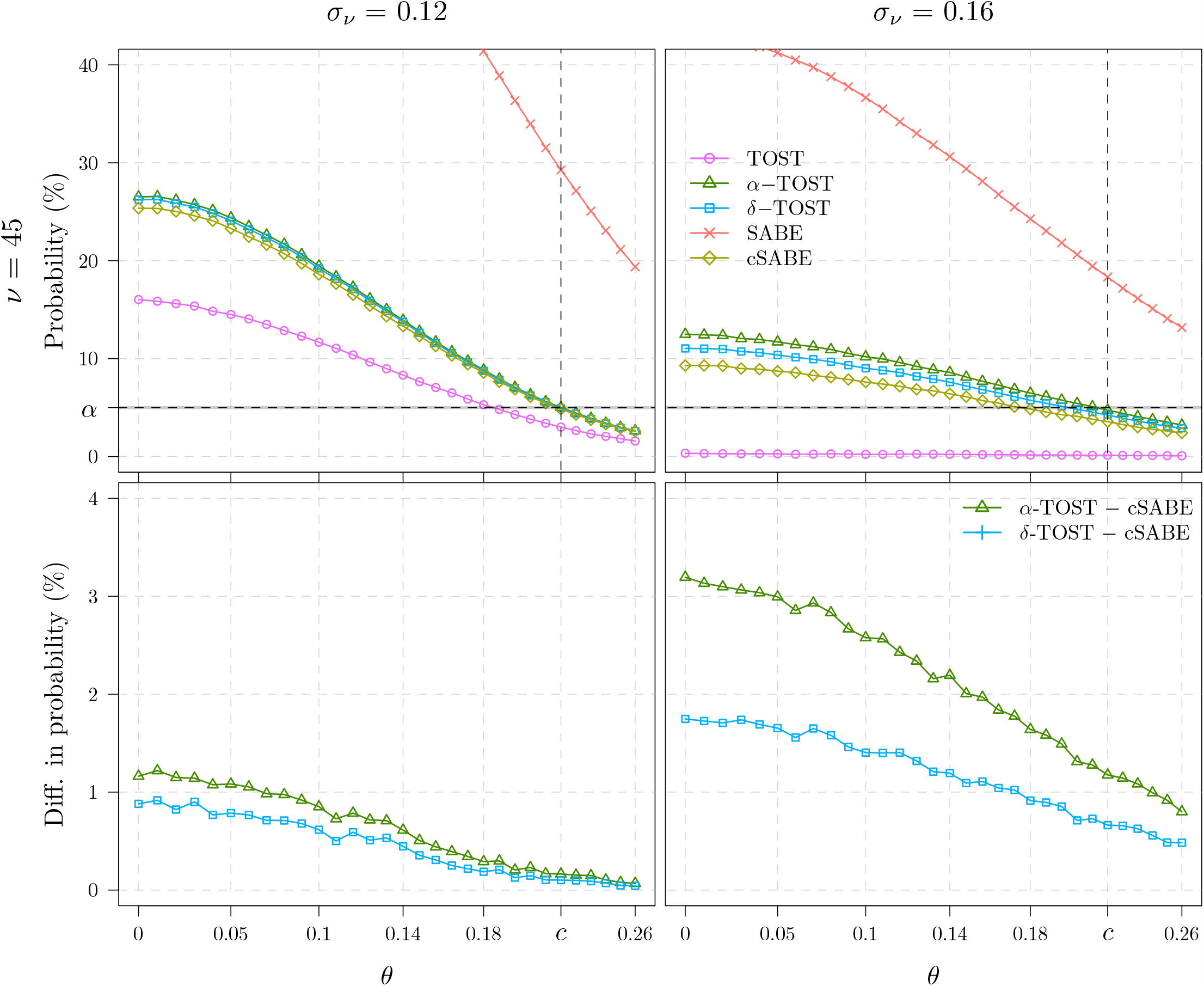
First row: empirical probability of declaring bioequivalence (*y*-axis) computed using the setting of Simulation 4, as a function of *θ* (*x*-axis) and *σ*_*ν*_ (columns), with *ν =*45, for the TOST (pink circles), the *α*-TOST (green triangles), the *δ*-TOST (blue squares), the SABE (red crosses) and the corrected SABE (light-green diamonds). The tight gray area stands for a 99% simulation error tolerance interval of (4.84, 5.16) corresponding to *α =*5% and *B =*10^5^ Monte Carlo samples. Second row: the difference of the empirical probabilities between the *α*-TOST and the cSABE (green triangles), and between the *δ*-TOST and the cSABE (blue triangles). Empirically, the TOST is quite conservative while the SABE is very liberal. In terms of power, the *α*-TOST uniformly dominates the other two methods and the *δ*-TOST uniformly dominates the cSABE.

**Figure 14:**
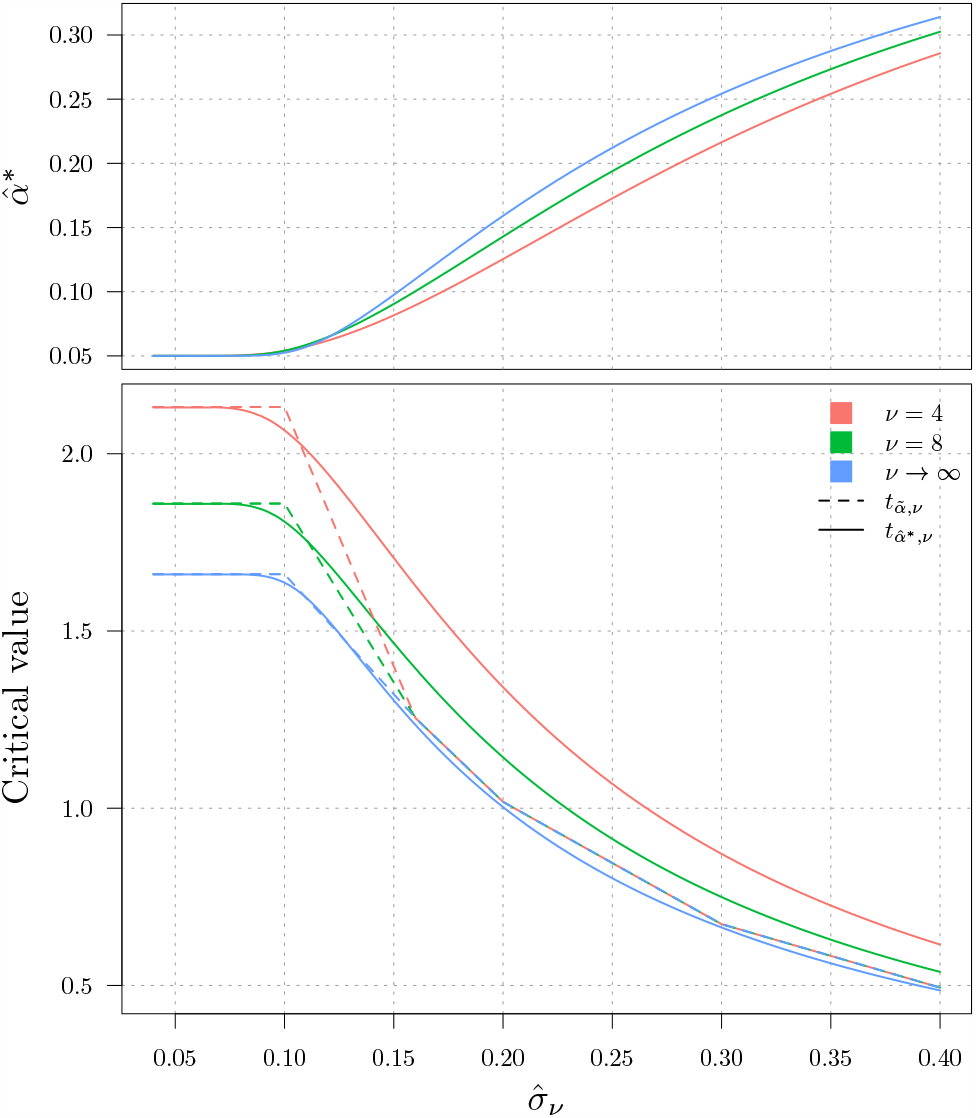
Upper panel: values of 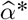 of the *α*-TOST (*y*-axis) as a function of 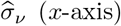 for different values of *ν* (coloured lines). Lower panel: comparison of the critical values (*y*-axis) obtained by the method of Cao and Mathew ^41^(dashed lines showing 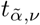) and of the *α*-TOST (solid lines showing 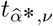 )as a function of 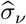 for different values of *ν* (coloured lines). Note that for all values of *ν* considered here and for values of 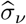above 0.1, the critical values of Cao and Mathew ^41^ correspond to a piecewise version of the critical values of the *α*-TOST obtained when *ν* is large. Therefore, their correction appears to be an approximation of the *α*-TOST, evaluated asymptotically, i.e., at *ν* → ∞.

### F Comparison of the *α*-TOST with Cao and Mathew’s Method

In Figure 14, the critical values for different values of *ν*, obtained by Cao and Mathew ^41^ (Table 1) and the ones obtained using the *α*-TOST, are compared. One can see that the method of Cao and Mathew ^41^ appears to be an approximation of the *α*-TOST, evaluated asymptotically, i.e., at *ν* → 8.

## Notes

### Competing Interest Statement

The authors have declared no competing interest.

### Summary of Updates

We have added short clarifications in three instances in the text. In page 9, starting with "This deviation from the nominal level...", in page 12 we have added the following sentence: "As expected, the power of all methods goes to one asymptotically as \sigma_\nu\to 0 (see Appendix B for details)" and page 22, we have simply replaced the definition of the asymptotic standard deviation by a definition of the asymptotic variance.

## References

[1] Metzler C. Bioavailability – A Problem in Equivalence. Journal of Pharmaceutical Sciences 1974; 30: 309–317.

[2] Westlake T. Symmetrical Confidence Intervals for Bioequivalence Trials. Biometrics 1976; 32: 741–744.

[3] Pallmann P, Jaki T. Simultaneous confidence regions for multivariate bioequivalence. Statistics in Medicine 2017; 36(29): 4585–4603.

[4] Senn S. Statistical Issues in Drug Development, 3rd edition. Wiley. 2021.

[5] Patterson S, Jones B. Bioequivalence and Statistics in Clinical Pharmacology. Boca Raton, FL: Chapman & Hall/CRC. 2006.

[6] Lakens D. Equivalence tests: A practical primer for t-tests, correlations, and meta-analyses. Social Psychological and Personality Science 2017; 8: 355–362.

[7] Sureshkumar H, Xu R, Erukulla N, Wadhwa A, Zhao L. “Snap on” or Not? A Validation on the Measurement Tool in a Virtual Reality Application. Journal of Digital Imaging 2022; 35(3): 692–703.

[8] O’Brien MW, Kimmerly DS. Is “not different” enough to conclude similar cardiovascular responses across sexes?. American Journal of Physiology-Heart and Circulatory Physiology 2022; 322: H355–H358.

[9] Wehrle FM, Bartal T, Adams M, et al. Similarities and Differences in the Neurodevelopmental Outcome of Children with Congenital Heart Disease and Children Born Very Preterm at School Entry. The Journal of Pediatrics 2022; 250: 29–37.e1.

[10] Sansone P, Giaccari LG, Aurilio C, et al. Comparative Efficacy of Tapentadol versus Tapentadol Plus Duloxetine in Patients with Chemotherapy-Induced Peripheral Neuropathy. Cancers 2022; 14: 4002.

[11] Branscheidt M, Ejaz N, Xu J, et al. No evidence for motor-recovery-related cortical connectivity changes after stroke using resting-state fMRI. Journal of Neurophysiology 2022; 127: 637–650.

[12] Feri F, Giannetti C, Guarnieri P. Risk-taking for others: An experiment on the role of moral discussion. Journal of Behavioral and Experimental Finance 2023; 37: 100735. doi: 10.1016/j.jbef.2022.100735

[13] Aggarwal M, Allen J, Coppock A, et al. A 2 million-person, campaign-wide field experiment shows how digital advertising affects voter turnout. Nature Human Behaviour 2023: 1–10.

[14] Meyners M. Equivalence tests – A review. Food Quality and Preference 2012; 26: 231–245.

[15] Lakens D, Scheel AM, Isager PM. Equivalence testing for psychological research: A tutorial. Advances in Methods and Practices in Psychological Science 2018; 1: 259–269.

[16] Mazzolari R, Porcelli S, Bishop DJ, Lakens D. Myths and methodologies: The use of equivalence and non-inferiority tests for interventional studies in exercise physiology and sport science. Experimental Physiology 2022; 107: 201–212.

[17] Wang K, Li Y, Chen B, et al. In Vitro Predictive Dissolution Test Should Be Developed and Recommended as a Bioequivalence Standard for the Immediate-Release Solid Oral Dosage Forms of the Highly Variable Mycopheno-late Mofetil. Molecular Pharmaceutics 2022; 19: 2048–2060.

[18] Schuirmann DJ. A comparison of the two one-sided tests procedure and the power approach for assessing the equivalence of average bioavailability. Journal of Pharmacokinetics and Biopharmaceutics 1987; 15(6): 657–680.

[19] Berger RL. Multiparameter hypothesis testing and acceptance sampling. Technometrics 1982; 24(4): 295–300.

[20] Munõz J, Alcaide D, Ocanã J. Consumer’s risk in the EMA and FDA regulatory approaches for bioequivalence in highly variable drugs. Statistics in Medicine 2016; 35: 1933–1943.

[21] Hsu JC, Hwang JTG, Liu HK, Ruberg SJ. Confidence intervals associated with tests for bioequivalence. Biometrika 1994; 81: 103–114.

[22] Berger RL, Hsu JC. Bioequivalence trials, intersection-union tests and equivalence confidence sets. Statistical Science 1996; 11: 283–319.

[23] Anderson S, Hauck WW. A new procedure for testing equivalence in comparative bioavailability and other clinical trials. Communications in Statistics A 1983; 12: 2663–2692.

[24] Brown HJTG, Munk A. An unbiased test for the bioequivalence problem. The Annals of Statistics 1997: 2345–2367.

[25] Liu JP, Chow SC. Bioequivalence trials, intersection-union tests and equivalence confidence set: Comment. Statistical Science 1996; 11: 306–312.

[26] Schütz H, Labes D, Wolfsegger MJ. Critical Remarks on Reference-Scaled Average Bioequivalence. Journal of Pharmacy & Pharmaceutical Sciences 2022; 25: 285–296.

[27] Guideline on the investigation of bioequivalence. European Medicines Agency. 2010. https://www.ema.europa.eu/en/documents/scientific-guideline/guideline-investigation-bioequivalence-rev1_en.pdf. Accessed July 10, 2023.

[28] Labes D, Schütz H. Inflation of Type I Error in the Evaluation of Scaled Average Bioequivalence, and a Method for its Control. Pharmaceutical Research 2016; 33: 2805–2814.

[29] Tothfalusi L, Endrenyi L, Arieta AG. Evaluation of bioequivalence for highly variable drugs with scaled average bioequivalence. Clinical pharmacokinetics 2009; 48: 725–743.

[30] Davit BM, Chen ML, Conner DP, et al. Implementation of a reference-scaled average bioequivalence approach for highly variable generic drug products by the US Food and Drug Administration. American Association of Pharmaceutical Scientists Journal 2012; 14: 915–924.

[31] Wonnemann M, Frömke C, Koch A. Inflation of the Type I Error: Investigations on Regulatory Recommendations for Bioequivalence of Highly Variable Drugs. Pharmaceutical Research 2015; 32: 135–143.

[32] Endrenyi L, Tothfalusi L. Bioequivalence for highly variable drugs: regulatory agreements, disagreements, and harmonization.. Journal of Pharmacokinetics Pharmacodynamics 2019; 46: 117–126.

[33] Molins E, Labes D, Schütz H, Cobo E, Ocanã J. An iterative method to protect the type I error rate in bioequivalence studies under two-stage adaptive 2×2 crossover designs. Biometrical Journal 2021; 63: 122–133.

[34] Tothfalusi L, Endrenyi L. An Exact Procedure for the Evaluation of Reference-Scaled Average Bioequivalence. American Association of Pharmaceutical Scientists Journal 2016; 18: 476–489.

[35] Ocanã J, Munõz J. Controlling type I error in the reference-scaled bioequivalence evaluation of highly variable drugs. Pharmaceutical Statistics 2019; 18: 583–599.

[36] Deng Y, Zhou XH. Methods to control the empirical type I error rate in average bioequivalence tests for highly variable drugs. Statistical Methods in Medical Research 2020; 29: 1650–1667.

[37] Phillips K. Power of the Two One-Sided Tests Procedure in Bioequivalence. Journal of Pharmacokinetics and Biopharmaceutics 1990; 18: 137–144.

[38] Owen DB. A special case of a bivariate non-central t-distribution. Biometrika 1965; 52: 437–446.

[39] Lehmann EL. Testing Statistical Hypothesis, 2nd edition. New York: Wiley. 1986.

[40] Palmes C, Bluhmki T, Funke B, Bluhmki E. Asymptotic properties of the two one-sided t-tests - new insights and the Schuirmann-constant. The International Journal of Biostatistics 2022; 18: 19–38.

[41] Cao L, Mathew T. A Simple Numerical Approach Towards Improving the Two One-Sided Test for Average Bioequivalence. Biometrical Journal 2008; 50: 205–211.

[42] Quartier J, Capony N, Lapteva M, Kalia YN. Cutaneous Biodistribution: A High-Resolution Methodology to Assess Bioequivalence in Topical Skin Delivery. Pharmaceutics 2019; 11: 484.

[43] Committee for Medicinal Products for Human Use. Draft Guideline on Quality and Equivalence of Topical Products. 2018. https://www.ema.europa.eu/en/documents/scientific-guideline/draft-guideline-quality-equivalence-topical-products_en.pdf. Accessed July 10, 2023.

[44] Jones B, Kenward MG. Design and Analysis of Cross-over Trials. CRC press. 2014.

[45] Dunnett CW, Gent M. Significance testing to establish equivalence between treatments, with special reference to data in the form of 2×2 tables. Biometrics 1977; 33: 593–602.

[46] Tu D. On the use of the ratio or the odds ratio of cure rates in establishing therapeutic equivalence of non-systemic drugs with binary clinical endpoints. Journal of Biopharmaceutical Statistics 1998; 8: 263–282.

[47] Schouten H, Kester A. A simple analysis of a simple crossover trial with a dichotomous outcome measure. Statistics in Medicine 2010; 29: 193–198.

[48] Lui KJ, Chang KC. Test non-inferiority (and equivalence) based on the odds ratio under a simple crossover trial. Statistics in Medicine 2011; 30: 1230–1242.

[49] Ostrovski V. Testing equivalence to binary generalized linear models with application to logistic regression. Statistics & Probability Letters 2022; 191: 109658.

[50] Rudin W. Principles of mathematical analysis, second edition. 1976.

[51] Vaart V. dAW. Asymptotic statistics. 3. Cambridge university press. 2000.

[52] Federer H. Geometric Measure Theory. Springer. 2014.

